# Copine-6 integrates calcium, phosphoinositide and Rab11 signals to coordinate glutamate receptor recycling

**DOI:** 10.64898/2026.08.26.747421

**Authors:** Jing Zhi Anson Tan, Ana Batallas-Borja, Mintu Chandra, Thanh Binh Nguyen, Se Eun Jang, Guoyang Gu, Lingrui Zhang, Kai-En Chen, Saroja Weeratunga, David B. Ascher, Jocelyn Widagdo, Brett M. Collins, Victor Anggono

## Abstract

Endosomal trafficking is a major pathway that delivers cell-surface proteins, including glutamate receptors, to support neurotransmission and normal brain functions. Activity-dependent insertion of glutamate receptors is essential for synaptic plasticity, learning and memory. Copine-6 is a neuronal-specific calcium (Ca^2+^) binding protein that mediates activity-induced exocytosis of α-amino-3-hydroxy-5-methyl-4-isoxazole propionic acid (AMPA)-type glutamate receptors. The activation of *N*-methyl-*D*-aspartate (NMDA) receptors triggers Ca^2+^-dependent translocation of Copine-6 to intracellular endosomal compartments. However, the mechanisms underlying the activity-dependent accumulation of Copine-6 in endosomes remain unknown. Here, we show that Copine-6 exhibits Ca^2+^-dependent binding to phosphatidylinositol-3-phosphate (PI(3)P) through the C2B domain and displays enhanced interaction with active Rab11a in a Ca^2+^-independent manner via the vWA domain. Mutations in the C2B that inhibit binding to PI(3)P not only block the activity-induced translocation of Copine-6 to early endosomes, but it also causes an aberrant accumulation of Copine-6 in recycling endosomes. Consequently, loss of Copine-6 expression impairs the efficient coupling of early and recycling endosomes and blocks activity-dependent delivery of both AMPA and NMDA receptors onto the neuronal plasma membrane. These defects can be restored by re-expressing wild-type Copine-6, but not the C2B phospholipid-binding mutant. Together, our findings establish Copine-6 as a molecular bridge that enhances coupling between the early and recycling endosomal membranes, thereby facilitating the activity-dependent forward trafficking of glutamate receptors to the neuronal plasma membrane to maintain synaptic potentiation.

**Significance Statement:** Activity-dependent trafficking of glutamate receptors is essential for synaptic plasticity, learning, and memory, but the mechanisms coordinating receptor transport through endosomal compartments remain unclear. This study identifies the neuronal calcium-binding protein Copine-6 as a molecular bridge that couples early and recycling endosomes during receptor trafficking. Copine-6 integrates calcium-dependent phospholipid binding and Rab11-mediated endosomal interactions to promote activity-dependent delivery of both AMPA and NMDA receptors to the neuronal surface. Loss of Copine-6 or disruption of its phospholipid-binding activity uncouples endosomal trafficking and impairs receptor insertion during synaptic potentiation. These findings reveal a key mechanism linking neuronal activity, endosomal organization, and glutamate receptor delivery to support synaptic function.

## Introduction

Dynamic modulation of synaptic strength within neuronal networks, a process termed synaptic plasticity, has long been regarded as the cellular basis of learning and memory (1–5). One of the most studied forms of synaptic plasticity is the *N*-methyl-*D*-aspartate receptor (NMDAR)-dependent long-term potentiation (LTP). During LTP, the activation of NMDARs triggers intracellular calcium (Ca^2+^)-dependent signaling cascades that drive a persistent increase in the number of α-amino-3-hydroxy-5-methyl-4-isoxazole propionic acid (AMPA)-type glutamate receptors (AMPARs) at the synapses, thereby enhancing excitatory neurotransmission (6–8). Rapid recruitment of synaptic AMPARs during early-LTP occurs through the lateral movement of extrasynaptic receptors on the plasma membrane (9, 10). To maintain LTP, the pool of extrasynaptic AMPARs is continuously replenished by activity-dependent recycling and exocytosis of receptors from the intracellular compartments to the neuronal plasma membrane (9, 11–14).

Coordinated membrane trafficking of AMPARs through the intracellular endosomal system dynamically controls their surface expression and synaptic strength (8, 15). Internalized AMPARs are sorted from early endosomes to late endosomes for degradation, or back to the plasma membrane via recycling endosomes (16–19). Differential lipid composition in distinct subcellular compartments, including the endosomes, is critical for the distribution and trafficking of membrane-associated proteins (20). Phosphoinositides (PIs) are a group of signaling phospholipids that regulate various aspects of AMPAR membrane trafficking in neurons. Phosphatidylinositol-(3,4,5)-trisphosphate (PIP_3_) and phosphatidylinositol-(4)-monophosphate (PI(4)P) mark the sites for AMPAR exocytosis on the plasma membrane and are essential for LTP (21–24). In contrast, the synthesis of phosphatidylinositol-(4,5)-bisphosphate (PI(4,5)P_2_) or phosphatidylinositol-(3,5)-bisphosphate (PI(3,5)P_2_) drives AMPAR internalization and contributes to synaptic depression (25, 26). Interestingly, LTP-inducing stimuli promote the production of phosphatidylinositol-(3)-phosphate (PI(3)P) on the early endosomes in the dendrites, which in turn supports the recycling of AMPARs via the sorting nexin 27 (SNX27)-retromer complex (27).

In addition to PIs, the identity and function of intracellular endosomal compartments are also defined by Rab small GTPases (20, 28), which ensure the proper trafficking and delivery of membrane-bound cargo proteins across distinct endosomal compartments. It is well established that Rab11-enriched recycling endosomes are the main source that supplies AMPARs to the neuronal plasma membrane during LTP (11, 29, 30). The coupling of Rab4-positive early endosomes with Rab11-containing recycling endosomes in the dendrite is controlled by a neuron-specific Rab4 effector GRASP-1 (31). The loss of GRASP-1 function impairs the incorporation of synaptic AMPARs during LTP, learning and memory (31, 32). Despite the physiological importance of endosomal coupling *in vivo*, how this process is regulated by Ca^2+^-dependent signaling during synaptic potentiation remains an open question.

Copine-6 is a neuron-specific postsynaptic Ca^2+^- and phospholipid-binding protein that mediates the activity-dependent exocytosis of AMPARs during synaptic potentiation (33). Loss of Copine-6 function impairs LTP and leads to deficits in learning and memory (34, 35). Copine-6 contains two N-terminal C2 domains and a C-terminal von Willebrand factor A (vWA) domain (33) 1A). The C2 domains bind Ca^2+^, which in turn triggers their interaction with phosphatidylserine (PS), thereby mediating Copine-6 translocation from the cytoplasm to the plasma membrane (33–38). In addition, Copine-6 protein is distributed across early, recycling and late endosomal compartments, with the highest abundance in Rab11-positive recycling endosomes (33). Importantly, Ca^2+^ binding also enhances Copine-6 localization in the recycling endosomes following synaptic potentiation, suggesting that Copine-6 may serve as a molecular link between Ca^2+^ signal and endosomal membrane trafficking in the dendrites. In this study, we investigated the molecular mechanisms underlying Copine-6 localization in endosomes and its functional relevance in regulating the delivery of surface glutamate receptors during synaptic potentiation.

## Results

### Copine-6 binds PIs in a Ca^2+^-dependent manner

The C2 domain generally binds PS and PIs in a Ca^2+^-dependent manner (39). Copine-6 has previously been reported to bind PS in the presence of Ca^2+^ (36). However, its interaction with other phospholipids is currently unknown. To address this question, we performed a qualitative lipid-binding assay by incubating a membrane spotted with an array of phospholipids with purified recombinant GST-Copine-6-C2AB (amino acids 1-305) proteins (Fig. 1B). As expected, Copine-6 exhibited the strongest binding to PS among fifteen phospholipids tested. We also observed weaker interactions between Copine-6 and the three phosphatidylinositol monophosphates (PIPs), namely PI(3)P, PI(4)P and PI(5)P (Fig. 1B).

**Figure 1.**
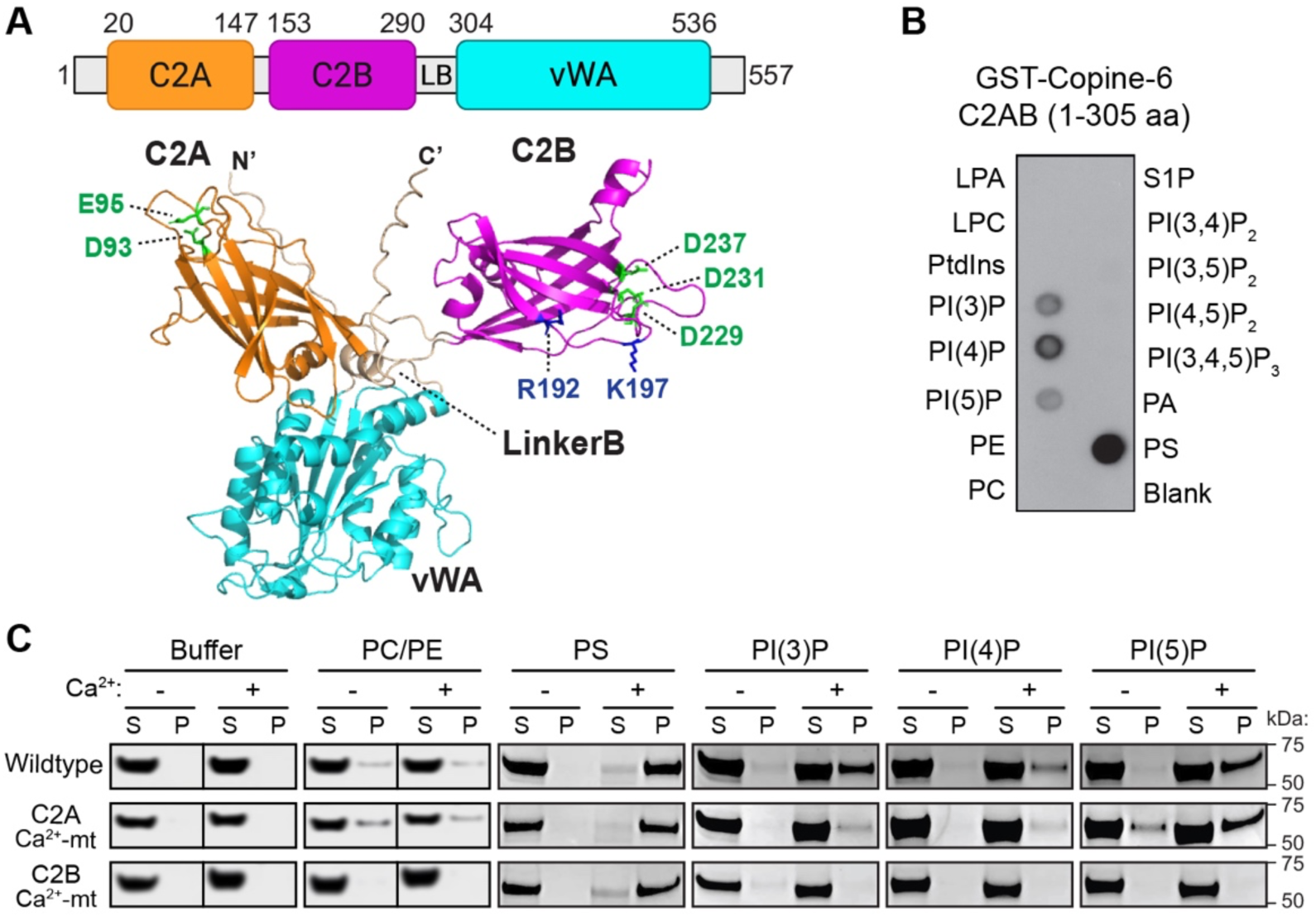
Copine-6 binds to phospholipids in a calcium-dependent manner. (A) Copine-6 domain structure (top) and the AlphaFold predicted structure (bottom) depicting the N-terminal C2A (orange) and C2B (magenta) domains and the vWA domain (cyan). The Ca^2+^-binding residues in C2A (Asp-93 and Glu-95) and C2B (Asp-229, Asp-231 and Asp-237) domains appear in green. The predicted phospholipid-binding residues in the C2B domain (Arg-192 and Lys-197) appear in blue. (B) A nitrocellulose membrane containing spots (100 pmole) of lysophosphatidic acid (LPA), lysophosphatidylcholine (LPC), phosphatidylinositol (PtdIns), phosphatidylinositol monophosphates [PI(3)P, PI(4)P and PI(5)P], phosphatidylethanolamine (PE), phosphatidylcholine (PC), sphingosine 1-phosphate (S1P), phosphatidylinositol bisphosphates [PI(3,4)P_2_, PI(3,5)P_2_ and PI(4,5)P_2_], phosphatidylinositol trisphosphates [PI(3,4,5)P_3_], phosphatidic acid (PA), phosphatidylserine (PS) and blank (control) was incubated with 0.5 μg/mL of purified GST-Copine-6-C2AB (1-305 amino acids) fusion protein. The binding was detected with anti-GST antibodies. (C) Purified recombinant His-Copine-6 full-length proteins (1-557 amino acids), either wildtype, C2A (D93N/E95A; C2A Ca^2+^-mt) or C2B (D229/231/237N; C2B Ca^2+^-mt) Ca^2+^-binding mutants were incubated with liposomes in the presence or absence of 2 mM Ca^2+^, followed by ultracentrifugation. The liposomes contain 0% (PC/PE), 30% PS or 10% PIPs. The unbound supernatant (S) and liposome-bound pellet (P) fractions were subjected to SDS-PAGE and Coomassie staining.

Next, we carried out liposome pelleting assays by incubating purified full-length Copine-6 recombinant proteins with phosphatidylcholine (PC) and phosphatidylethanolamine (PE) liposomes containing 30 mol% PS or 10 mol% PIPs. Wild-type Copine-6 proteins strongly bound PS liposomes only in the presence of Ca^2+^ and were absent in the pellet fraction when incubated with PC/PE only liposomes (Fig. 1C). Surprisingly, mutations of acidic residues that coordinate Ca^2+^ ions in the C2A (D93N/E95A; C2A Ca^2+^-mt) or C2B domain (D229/231/237N; C2B Ca^2+^-mt) did not affect PS liposome binding to full-length Copine-6 (Fig. 1C). However, we observed a partial reduction in PS binding to recombinant Copine-6-C2AB proteins when all five acidic residues (D93N/E95A and D229/231/237N) were mutated (SI Appendix, Fig. S1), suggesting a potential cooperative PS-binding by the two C2 domains.

Consistent with the lipid-strip data, we also observed robust Ca^2+^-dependent binding of wild-type Copine-6 proteins to PI(3)P, PI(4)P and PI(5)P liposomes (Fig. 1C). C2B Ca^2+^-mt failed to bind all three PIPs (Fig. 1C). In contrast, the C2A Ca^2+^-mt, which weakens the binding of Copine-6 to Ca^2+^ (33), only had mild effects on PIP binding (Fig. 1C). No interaction with any of the three PIPs was observed with Copine-6-C2AB double mutants (SI Appendix, Fig. S1). Taken together, these results demonstrate that, in addition to PS, Copine-6 directly interacts with PIPs upon Ca^2+^ binding, likely through the C2B domain.

### Copine-6 binds PIPs via the C2B domain

To further characterize the lipid-binding properties of Copine-6, we performed multiple sequence alignments of its C2 domains with those of protein kinase Cα (PKCα), synaptotagmin-1 (Syt-1) and Rabphilin-3A (Rph3a), and identified two conserved positively charged residues in the C2A (Arg-59 and Arg-64) and C2B (Arg-192 and Lys-197), which are predicted to interact with phospholipids (Fig. 2A). Structural alignment of the AlphaFold-predicted Copine-6 C2B domain with the C2 domain of PKCα revealed positional conservation of Arg-192 and Lys-197 to Lys-211 and Arg-216 in the C2 domain of PKCα known to directly interacts with PI(4,5)P_2_ and PS, respectively (Fig. 2B) (40). The presence of a PI binding pocket in the C2B domain is further supported by surrounding key tyrosine residues (Tyr-179, Tyr-228 and Tyr-230) that contribute to the phospholipid-binding interface (Fig. 2C). Unlike the C2B domain, we noted that despite the structural alignment of Arg-59 in the C2A domain with Arg-192 (Copine-6 C2B domain) and Lys-211 (PKCα C2 domain), the presence of Pro-101 creates a steric hindrance, preventing PI binding to the C2A domain. In addition, the corresponding tyrosine residues that coordinate phospholipid binding are absent in the C2A domain (Fig. 2C), further diminishing the propensity of Copine-6 C2A domain to bind PIs.

**Figure 2.**
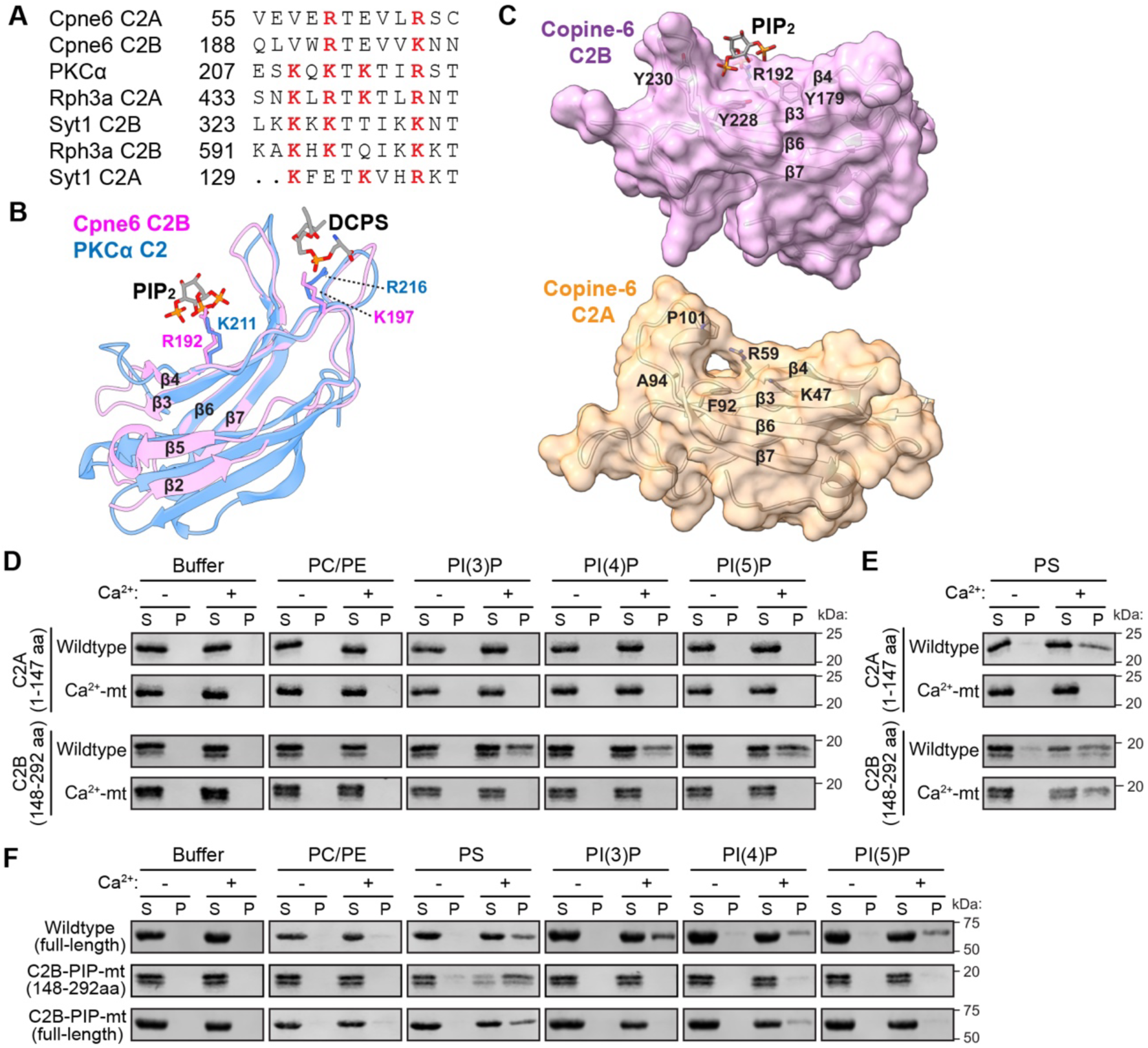
Copine-6 binds phosphatidylinositol monophosphates via the C2B domain. (A) Multiple sequence alignment of the â4 strand within the C2 domain of Copine-6 (Cpne6), protein kinase C-alpha (PKCá), rabphilin-3a (Rph3a) and synaptotagmin-1 (Syt1). The conserved arginine (R) and lysine (K) residues involved in phospholipid-binding are indicated in red. (B) Structural comparison of the AlphaFold-predicted C2B domain of Copine-6 (pink) with the C2 domain of PKCα (blue) bound to PI(4,5)P_2_ (PIP_2_) and dicaproyl phosphatidylserine (DCPS) [PDB: 1DSY and 3GPE]. Arg-192 and Lys-197 in Copine-6 C2B domain shows positional conservation with Lys-211 and Arg-216 in PKCα known to interact with PIP_2_ and DCPS. (C) Surface representations of AlphaFold structures of Copine-6 C2B (top, pink) and C2A (bottom, golden) domains. The homologous PIP_2_ binding pocket in C2B is occluded in the C2A domain due to the presence of atomic densities formed by Pro-101 and Arg-59. (D–F) Liposome pelleting assays. Purified recombinant His-Copine-6-C2A (amino acids 1-147) or His-Copine-6-C2B (amino acids 148-292), either wildtype (WT) or the Ca^2+^-binding mutants (Ca^2+^-mt) were incubated with liposomes containing PIPs (D) or PS (E) in the presence or absence of 2 mM Ca^2+^, followed by ultracentrifugation. (F) Full-length His-Copine-6, either WT or the phospholipid-binding deficient mutant (R192Q/K197A; C2B PIP-mt) and His-Copine-6-C2B PIP-mt were incubated with liposomes containing 10% PIPs were subjected to the same assay. The unbound supernatant (S) and liposome-bound pellet (P) fractions were subjected to SDS-PAGE and Coomassie staining.

To test this prediction, we performed liposome-binding assays using purified recombinant proteins of the individual C2A (amino acids 1-147) and C2B (amino acids 148-292) domains. As predicted, the C2B, but not the C2A domain, bound PI(3)P, PI(4)P and PI(5)P in a Ca^2+^-dependent manner (Fig. 2D). None of these PIPs interacted with the C2B Ca^2+^-mt, confirming that the Ca^2+^-dependent binding between Copine-6 and PIPs is mediated by the C2B domain (Fig. 2D). The individual Copine-6 C2 domains were not found in the pellet fraction when incubated with the buffer only or PC/PE liposomes, demonstrating the specificity of our assays (Fig. 2D).

We also investigated the ability of individual Copine-6 C2 domains to bind PS using the same liposome pelleting assay. As expected, both wild-type C2A and C2B proteins were detected in the pellet fraction when Ca^2+^ was present in the reaction (Fig. 2E). Mutations of the Ca^2+^ binding sites in the C2A domain completely abolished its interaction with PS (Fig. 2E). Consistent with the findings obtained with full-length Copine-6, the Ca^2+^ binding mutant in the isolated C2B domain also did not affect Ca^2+^-induced PS binding (Fig. 2E). To gain further insights, we performed molecular dynamics (MD) simulations of the C2B domain in the presence of Ca^2+^ ions, PS and PI(3)P headgroups. It revealed that the PS headgroup (hPS) stably occupied the Ca^2+^-binding site beyond Asp-229, Asp-231, and Asp-237, and included Asp-167 and Asp-173 (SI Appendix, Fig. S2A and Video 1). These two additional aspartates have also been previously shown to affect Copine-6 association with the plasma membrane when mutated (35, 38). Although hPS was no longer stabilized around Asp-173, Asp-229, Asp-231, and Asp-237 in the C2B Ca^2+^-mt, it exhibited preferential interaction with Asp-167 (SI Appendix, Fig. S2B and Video 2). This may explain the apparent residual binding of PS to Copine-6 C2B Ca^2+^-mt.

Unlike hPS, the PI(3)P headgroup (hPI3P) was predominantly localized near Arg-192 for part of the simulation (SI Appendix, Fig. S3A and Video 1). We also found that Lys-197 was the closest positively charged residue to interact with hPS (SI Appendix, Fig. S3B and Video 1). Based on these observations and the structural alignment of Arg-192 and Lys-197 with the known PKCα binding sites to PI(4,5)P_2_ and PS, we mutated these two positively charged residues in the C2B domain (R192Q and K197A) and tested their effects on Ca^2+^-induced phospholipid binding to Copine-6. As predicted, the C2B R192Q/K197A mutant abolished Copine-6 binding to PI(3)P, PI(5)P, and reduced binding to PI(4)P in the presence of Ca^2+^ (Fig. 2F). However, PS-binding was unaffected by the C2B R192Q/K197A mutant in the liposome binding assay (Fig. 2F). This is perhaps not surprising considering the distance between Lys-197 and hPS was greater than 5 Å as determined from MD simulations (SI Appendix, Fig. S3B and Video 1). The same results were obtained when these mutations were introduced into full-length Copine-6 (Fig. 2F). We therefore isolated a Copine-6 mutant that severely weakens its specific binding to mono-phosphorylated PIs, but not to PS. We refer to the R192Q/K197A mutant as the C2B PIP-binding mutant (PIP-mt) hereafter.

### PI(3)P binding mediates activity-induced translocation of Copine-6 to early endosomes

The differential distribution of PIPs across distinct endosomal compartments is essential for regulating intracellular membrane trafficking (20). During LTP, the syntheses of PI(3)P and PI(4)P are elevated and support an increase in endosomal recycling and delivery of AMPARs to the dendritic plasma membrane (24, 27). We have previously shown that Copine-6 associates with intracellular endosomal compartments and becomes enriched in the Rab11-positive recycling endosome during synaptic potentiation (33). However, whether this process is mediated by the direct binding of Copine-6 with PIPs remains unknown. To address this question, we expressed Copine-6-GFP (wild-type or the C2B PIP-mt) with FYVE-mCherry, which marked the PI(3)P-enriched intracellular membrane compartments (primarily early endosomes) in cultured hippocampal neurons (Fig. 3A). Neurons were stimulated with glycine to induce chemical-LTP (cLTP) and fixed. Although there was no significant difference in the distribution of wild-type Copine-6-GFP and the C2B PIP-mt under basal conditions, glycine-induced accumulation of Copine-6 in the FYVE-positive endosomes was inhibited in the soma (Fig. 3A and B) and dendrites (Fig. 3C and D) of neurons expressing Copine-6-GFP C2B PIP-mt. The activity-dependent enrichment of Copine-6-GFP in the PI(3)P-containing membrane compartment correlated with an increase in the number of FYVE-mCherry-positive puncta in the dendrites (Fig. 3C and E). Importantly, glycine-induced synthesis of PI(3)P remained intact in neurons expressing the Copine-6-GFP C2B PIP-mt (Fig. 3C and E), indicating that the impairment in activity-induced accumulation of the mutant Copine-6 to PI(3)P-positive endosomes was not due to the loss of PI(3)P synthesis. Collectively, these data demonstrate that Ca^2+^-dependent binding of Copine-6 to PI(3)P is essential for activity-induced translocation of Copine-6 to the early endosomes during synaptic potentiation.

**Figure 3.**
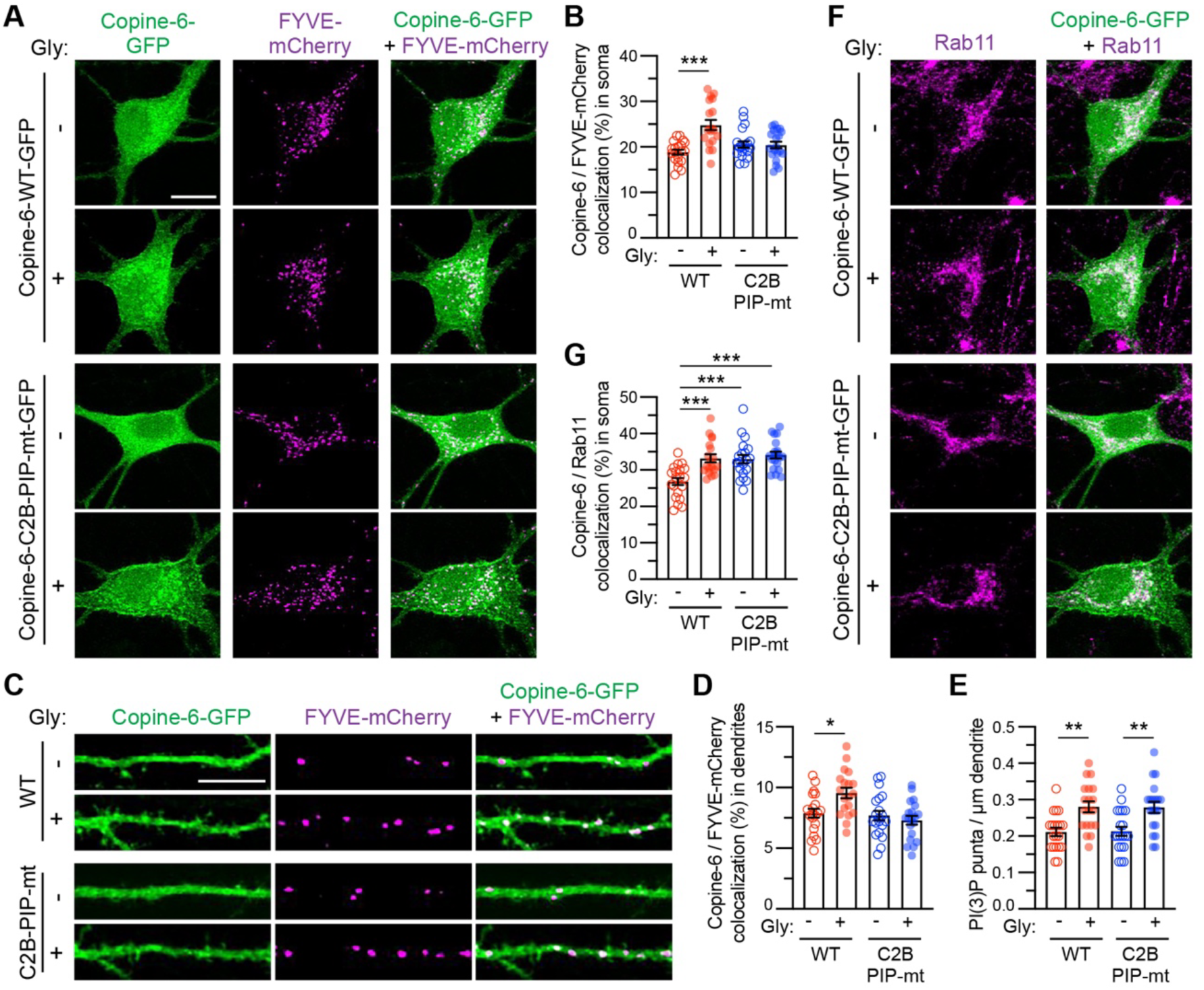
PI(3)P binding mediates activity-induced accumulation of Copine-6 to early endosomes. (A) Primary hippocampal neurons were co-transfected with plasmids encoding Copine-6-GFP (green), either wild-type (WT) or PIP-binding deficient mutant (C2B PIP-mt) and the PI(3)P reporter, FYVE-mCherry (magenta) at days *in vitro* (DIV) 12. At DIV15, neurons were stimulated with glycine for 5 min, fixed and visualized by confocal microscopy. Scale bar, 10 μm. (B) Quantification of Copine-6-GFP colocalization in the FYVE-mCherry-positive compartments using Mander’s coefficient. (C) Representative images of secondary dendrites of primary hippocampal neurons co-expressing Copine-6-GFP (WT or C2B PIP-mt, green) and FYVE-mCherry (magenta) under basal conditions or following 5 min of glycine stimulation. Scale bar, 10 μm. (D and E) Quantification of Copine-6-GFP colocalization with FYVE-mCherry using Mander’s coefficient (D) and the number of FYVE-mCherry-positive puncta in the first 30 μm of secondary dendrites (E). (F) In the same sets of experiments as (A), neurons were also immunostained for endogenous Rab11 (magenta). (G) Quantification of Copine-6-GFP colocalization in Rab11-positive endosomes using Mander’s coefficient. All data are represented as mean ± SEM. *n* = 19-20 neurons per group from 4 independent experiments. * *P* < 0.05, ** *P* < 0.01 and *** *P* < 0.001 using one-way ANOVA with Tukey’s multiple comparison test.

In the same experiments, we also determined the distribution of Copine-6-GFP in recycling endosomes by immunostaining with an antibody against endogenous Rab11 (Fig. 3F). As expected, glycine stimulation enhanced Copine-6-GFP localization in Rab11-positive recycling endosomes (Fig. 3F and G). In contrast, the C2B PIP-mt displayed a marked increase in the basal association with the recycling endosomes compared to wild-type Copine-6-GFP, reaching a level that was comparable to those in potentiated neurons (Fig. 3F and G). Indeed, glycine stimulation did not increase the level of Copine-6-GFP C2B PIP-mt in the recycling endosomes (Fig. 3F and G). These results indicate that the inability of Copine-6 to bind PI(3)P leads to an aberrant accumulation in the recycling endosomes.

### Copine-6 interacts with the Rab11 small GTPase protein

Next, we investigated the potential mechanism underlying the redistribution of Copine-6 C2B PIP-mt to Rab11-positive compartments. Recycling endosomes are particularly enriched in PI(4)P and are characterized by their association with Rab11 (20, 41). Copine-6 C2B PIP-mt exhibited a reduced binding to PI(4)P and therefore could not explain the apparent increase in its localization to recycling endosomes. Given that Copine-6 binds to the small GTPase Rac1 (35), we reasoned that it might also interact with Rab11. To test this hypothesis, we performed GST pull-down assays using lysates from HEK293T cells co-expressing wild-type GFP-Rab11a and GST-Copine-6. We found that GFP-Rab11a specifically interacted with GST-Copine-6 but not the GST itself (Fig. 4A). The interaction between Rab11a and Copine-6 C2AB (amino acids 1-305) was comparable to that of the full-length protein (Fig. 4A and B). On the other hand, the isolated Copine-6 vWA domain (amino acids 306-557) exhibited the highest level of binding to GFP-Rab11 (Fig. 4A and B). These data suggest that Rab11a recognizes two binding interfaces on Copine-6, with the vWA domain as the primary binding site. Interestingly, the binding of GFP-Rab11a to Copine-6 C2B Ca^2+^-mt was comparable to the wild-type protein (Fig. 4C and D), indicating that the binding to Rab11a is independent of Copine-6 Ca^2+^-binding. In contrast, Copine-6 C2B PIP-mt exhibited a significantly enhanced binding to Rab11a (Fig. 4C and D), which correlates with the increased Copine-6 localization in the recycling endosomes (Fig. 3F and G).

**Figure 4.**
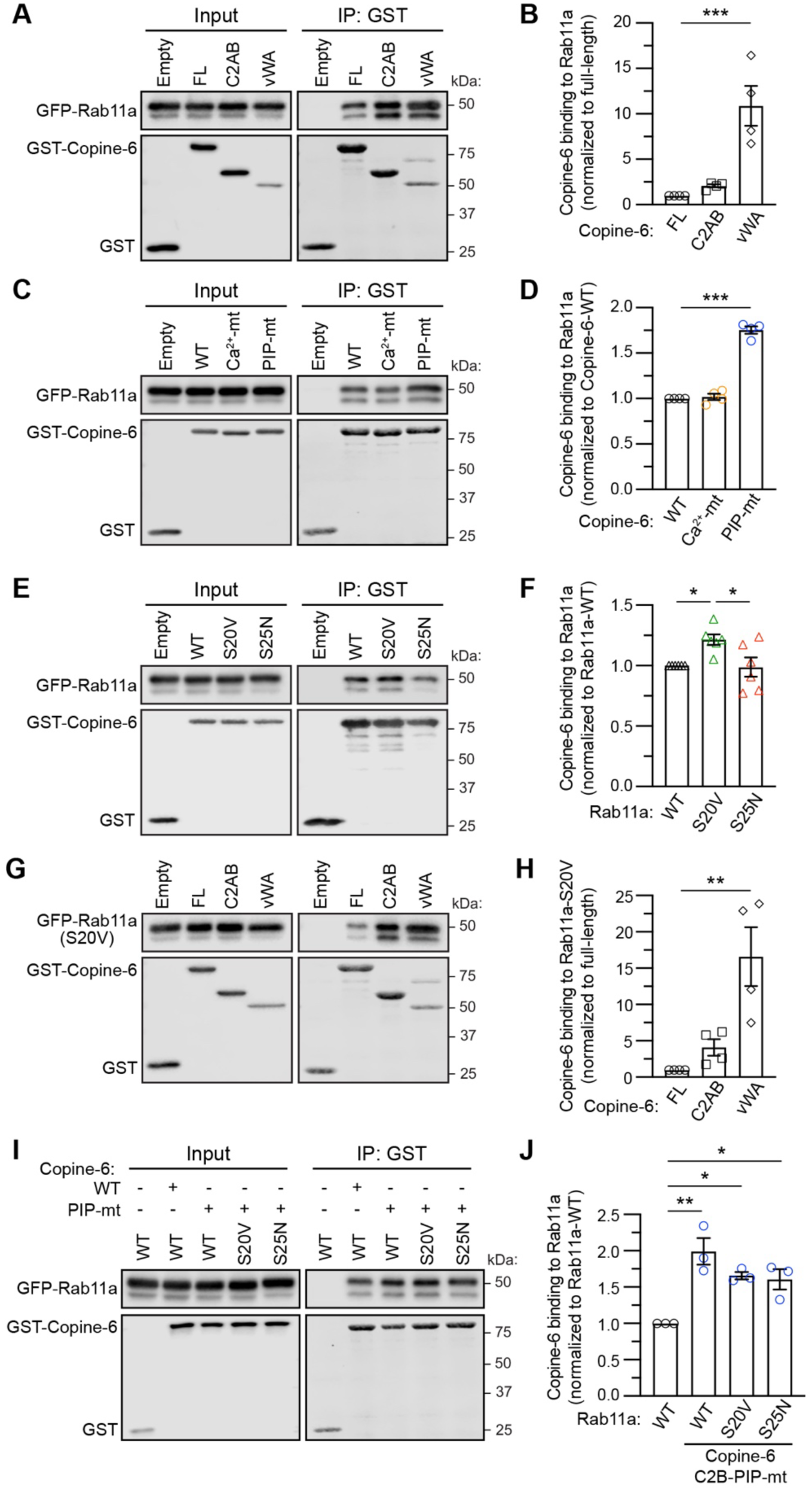
Copine-6 forms a complex with Rab11. (A) HEK293T cells were co-transfected with plasmids encoding GFP-Rab11a and GST-Copine-6, either full-length protein (amino acids 1-557), C2AB domain (amino acids 1-305) or the vWA domain (amino acids 306-557). GST alone serves as the control. Cells were lyzed and incubated with GSH-Sepharose beads. Total lysates (Input) and bound proteins (Pull-down) were resolved by SDS-PAGE and analyzed by western blotting with specific antibodies against GFP or GST. (B) Quantification of the relative binding of GFP-Rab11a binding to GST-Copine-6 proteins. (C and D) GST pull-down assays and quantification of GFP-Rab11a binding to GST-Copine-6 proteins, either wild-type (WT), Ca2+-binding deficient mutant (D229/231/237N; C2B Ca^2+^-mt) or the phospholipid-binding deficient mutant (R192Q/K197A; C2B PIP-mt). (E and F) GST pull-down assays and quantification of GST-Copine-6 binding to GFP-Rab11a proteins, either WT, the constitutive active (S20V) or the constitutive inactive (S25N) mutants. (G and H) GST pull-down assays and quantification of GFP-Rab11a S20V binding to GST-Copine-6 proteins, either full-length, C2AB or vWA domains. (I and J) GST pull-down assays and quantification of GST-Copine-6 C2B PIP-mut binding to GFP-Rab11a, either WT, S20V or S25N mutants. Data represent mean ± SEM from *N* = 3 to 6 independent experiments. * *P* < 0.05, ** *P* < 0.01 and *** *P* < 0.001 using one-way ANOVA with Tukey’s multiple comparison test.

The fact that Ca^2+^ binding to Copine-6 is not required for its interaction with Rab11a raises the possibility that activity-dependent translocation of Copine-6 to the recycling endosomes could depend on the state of nucleotide-bound Rab11. Using the same GST pull-down assays, we observed a small but significant increase in the binding of Copine-6 to the constitutively active GFP-Rab11a S20V (42) compared to wild-type or the S25N dominant negative mutant (Fig. 4E and F), demonstrating that Copine-6 is a downstream effector of active Rab11. We also confirmed that the constitutively active Rab11a S20V primarily interacts with Copine-6 via the vWA domain (Fig. 4G and H). Interestingly, this state-dependent binding was no longer observed in the Copine-6 C2B PIP-mt (Fig. 4I and J).

### Copine-6 mediates activity-induced coupling of early and recycling endosomes

The ability of Copine-6 to accumulate in early and recycling endosomes positions it as a critical molecule that mediates activity-dependent endosomal coupling in neurons during synaptic potentiation. To determine the role of Copine-6 in endosomal coupling, we assessed the co-localization of endogenous EEA1 (a marker of early endosomes) and GFP-Rab11a in control (Cas9-tdTomato) and Copine-6 knockdown neurons (Cas9-tdTomato + Copine-6 sgRNA). As expected, we observed a significant increase in the levels of EEA1/GFP-Rab11a co-localization in the soma (SI Appendix, Fig. S4A and B) and dendrites (Fig. 5A and B) of control primary hippocampal neurons upon glycine stimulation. Conversely, the activity-induced increase in EEA1/GFP-Rab11a co-localization was abolished in Copine-6 knockdown neurons (Fig. 5A and B and SI Appendix, Fig. S4A and B). This deficit could be fully restored by re-expressing the sgRNA-resistant wild-type Copine-6 but not the C2B PIP-mt (Fig. 5A and B and SI Appendix, Fig. S4A and B). Because recycling endosomes within the dendrites undergo dynamic structural changes from tubular to compact forms that provide membranes to support structural and functional plasticity of the spines (43, 44), we next measured the coefficient of variation of GFP-Rab11a distribution along the dendritic shaft as a proxy for the diffuseness of their fluorescence. We found that manipulating Copine-6 expression or introducing the PIP-binding-deficient mutant did not affect recycling endosome dynamics in dendrites (Fig. 5A and C). Together, our results demonstrate that Ca^2+^-dependent binding of Copine-6 to PI(3)P is an essential step that mediates activity-induced coupling between the early and recycling endosomes.

**Figure 5.**
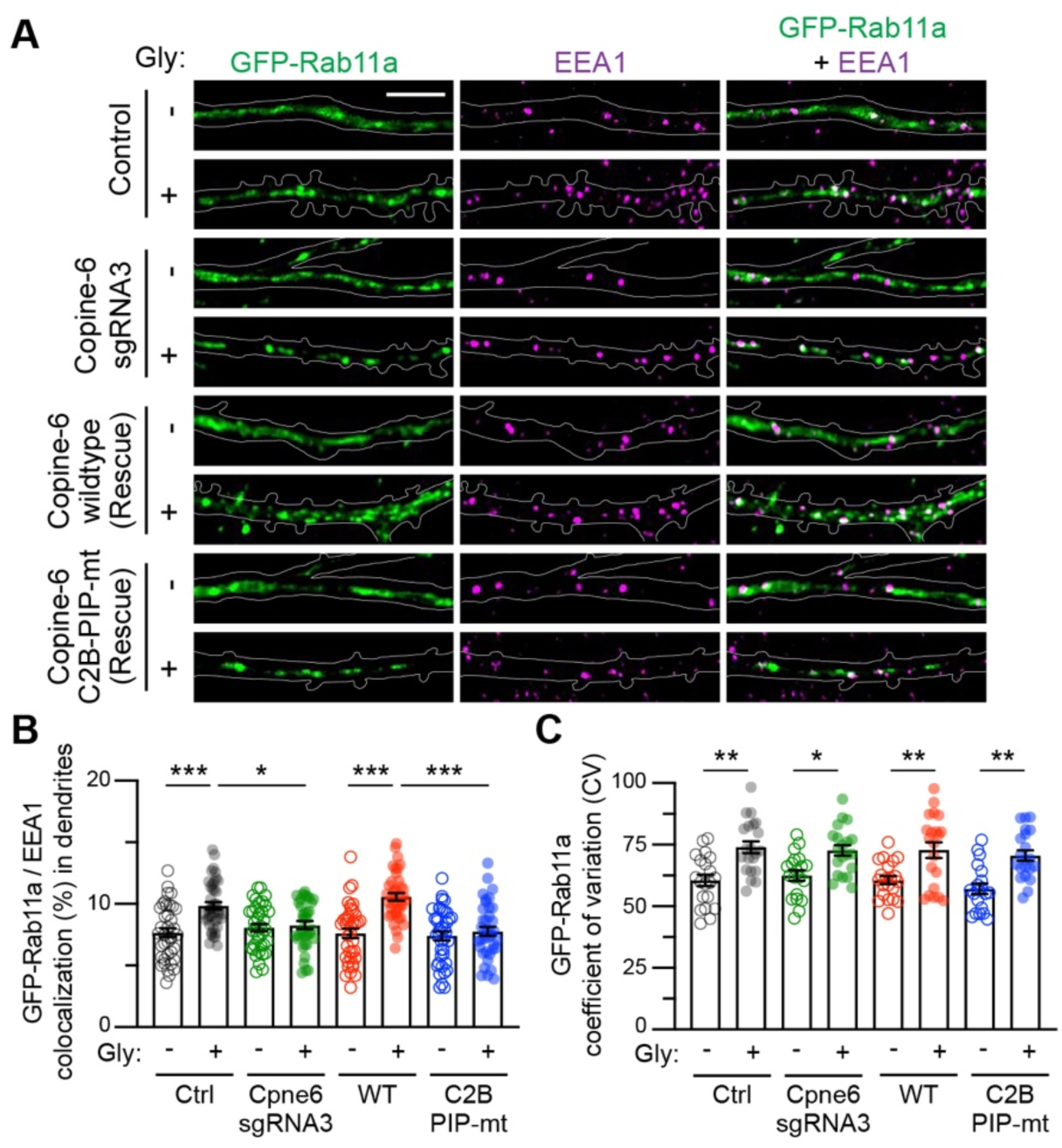
Copine-6 mediates activity-induced coupling of early and recycling endosomes. (A) The plasmid encoding GFP-Rab11a (green) was transfected into primary hippocampal neurons expressing Cas9-T2A-tdTomato only (control), Cas9-T2A-tdTomato-Copine-6 sgRNA3 (Cpne6 knockdown), Cas9-T2A-tdTomato-Copine-6 sgRNA3 plus sgRNA3-resistant Copine-6 wild-type (Rescue WT) or Cas9-T2A-tdTomato-Copine-6 sgRNA3 plus sgRNA3-resistant Copine-6 R192Q/K197A (Rescue C2B PIP-mt) at DIV 12. Neurons were stimulated with glycine for 5 min at DIV15, fixed and immunostained for endogenous EEA1 (magenta). Representative images of the dendritic segment of the co-transfected neurons are shown. Scale bars, 10 μm. (B) Quantification of dendritic the fraction of GFP-Rab11a colocalizing within EEA1-positive compartments using Mander’s coefficient. (C) The diffuseness of GFP-Rab11a along the dendrites was determined by calculating the coefficient of variance (CV) of GFP signals. Each data point represents the average CV of GFP-Rab11a from 4-5 secondary dendrites per neuron. Data represent mean ± SEM. *n* = 20-21 neurons per group from 3 independent experiments. * *P* < 0.05, ** *P* < 0.01 and *** *P* < 0.001 using one-way ANOVA with Tukey’s multiple comparison test.

### Copine-6 binding to PI(3)P is essential for activity-induced AMPAR exocytosis

The coupling of the early and recycling endosomes is critical for activity-induced AMPAR recycling and synaptic plasticity (11, 31). We have previously shown that Copine-6 and its interaction with Ca^2+^ are essential for activity-induced forward trafficking of AMPARs to the neuronal plasma membrane during synaptic potentiation (33). To interrogate the functional significance of Copine-6 binding to PI(3)P in regulating the surface expression of AMPARs to the plasma membrane, we first performed surface biotinylation assays in Copine-6-depleted neurons that re-expressed sgRNA-resistant myc-Copine-6, either wild-type or the C2B PIP-mt, under basal conditions or following glycine stimulation. As expected, neurons expressing wild-type myc-Copine-6 exhibited enhanced surface AMPARs upon glycine stimulation (Fig. 6A and B). In contrast, the expression of myc-Copine-6 C2B PIP-mt failed to restore the glycine-induced increase in the level of surface AMPARs without affecting the total protein abundance (Fig. 6A-C).

**Figure 6.**
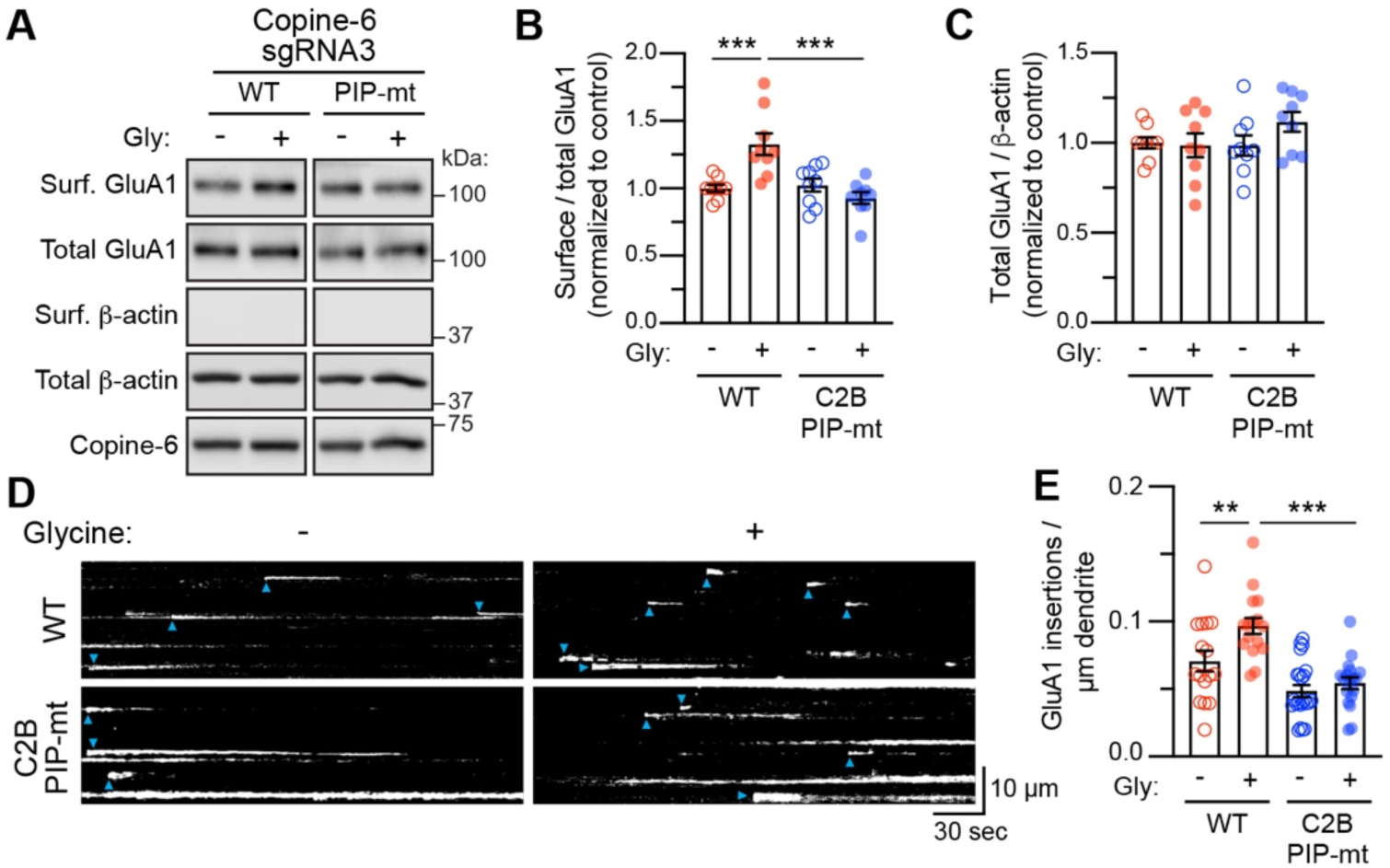
Copine-6 binding to PI(3)P is required for activity-induced AMPAR exocytosis. (A) Primary cortical neurons expressing Cas9-GFP-Copine-6 sgRNA3 (Copine-6 knockdown) were co-transduced with lentiviral particles expressing sgRNA3-resistant myc-Copine-6, either wild-type (WT) or R192Q/K197A (C2B PIP-mt) DIV9. At DIV15, transduced neurons were stimulated with glycine for 5 min and subjected to a surface biotinylation assay. The relative amounts of surface and total proteins were assessed by western blotting using specific antibodies against GluA1, Copine-6 and β-actin. (B and C) Quantification of the surface to total GluA1 ratio (B) and total GluA1 to β-actin ratio (C). Data were normalized to unstimulated myc-Copine-6 WT control neurons (*n* = 9 cultures per group from 7 independent experiments). (D) Primary hippocampal neurons were co-transfected with SEP-GluA1 reporter construct and Cas9-T2A-tdTomato-Copine-6 sgRNA3 with sgRNA3-resistant Copine-6 (WT or C2B PIP-mt). Plasma membrane insertion of SEP-GluA1 in dendrites were visualized on a total internal reflection fluorescence (TIFR) microscope over a 5 min period under basal conditions or following glycine stimulation. Representative *y*-axis by time (*y*-t) maximum intensity projections of SEP-GluA1 insertion events in a 30 μm segment of a secondary dendrite from a neuron in each group. Each GluA1 insertion event is marked by a blue arrowhead. (E) Quantification of the GluA1 insertion events per ìm of dendrite per min following glycine stimulation (*n* = 16-23 neurons per group from 3 independent experiments). All data are represented as mean ± SEM. \*\**P* < 0.01 and \*\*\**P* < 0.001 using one-way ANOVA with a Dunnett’s multiple comparison test.

Next, we directly monitored exocytosis of AMPARs by imaging super-ecliptic pH-sensitive-GFP (SEP)-tagged GluA1 in living hippocampal neurons using total internal reflection fluorescence (TIRF) microscopy (33). Using the same molecular replacement strategy, we observed a robust increase in the number of SEP-GluA1 insertions on the dendritic plasma membrane of neurons expressing wild-type myc-Copine-6 following glycine stimulation (Fig. 6D and E). However, glycine-induced elevation in SEP-GluA1 insertion was impaired in the hippocampal neurons expressing myc-Copine-6 C2B PIP-mt (Fig. 6D and E). To rule out that this deficit was not due to an impairment in the interaction between Copine-6 C2B PIP-mt with GluA1, we performed proximity-based ligation assays (PLA) and did not find any significant changes in the proximal binding between these two proteins under basal conditions or following glycine stimulations (SI Appendix, Fig. S5A-C). Our results demonstrate that Ca^2+^-dependent interaction between Copine-6 and PI(3)P facilitates the coupling of early and recycling endosomes, thereby promoting the recycling and surface delivery of AMPARs during synaptic potentiation.

### Copine-6 binding to PI(3)P is essential for activity-induced NMDAR exocytosis

AMPARs are not the only molecules mobilized from intracellular endosomes to the plasma membrane during LTP (45). GluN2A-containing NMDARs represent another member of the ionotropic glutamate receptor family that are rapidly inserted into the neuronal membrane following synaptic potentiation (46–51). First, we performed a pull-down assay and found that Copine-6 can be pulled down by GST-GluN2A C-terminal fusion protein (SI Appendix, Fig. S6A), suggesting that Copine-6 and GluN2A form a complex in cells. Given that activity-induced insertion of GluN2A-NMDARs relies on the endosomal recycling pathway (46), we explored whether Copine-6 might regulate this process by performing a series of surface biotinylation assays. We found that the glycine-induced increase in surface GluN2A-NMDARs was abolished in Copine-6 knockdown neurons using two independent sgRNAs (Fig. 7A and B). Loss of Copine-6 did not affect the steady-state expression of total and surface GluN2A-NMDARs in primary neurons (SI Appendix, Fig. S6B). Activity-induced surface delivery of GluN2A-NMDARs was restored in Copine-6 knockdown neurons by the re-expression of the wild-type, but not the Ca^2+^-(Fig. 7C and D) or PI(3)P-binding deficient Copine-6 mutants (Fig. 7E and F). Collectively, our data support a model in which Copine-6 regulates the Ca^2+^-dependent coupling of endosomal compartments for the sorting and trafficking of glutamate receptors to the plasma membrane during synaptic potentiation (Fig. 8).

**Figure 7.**
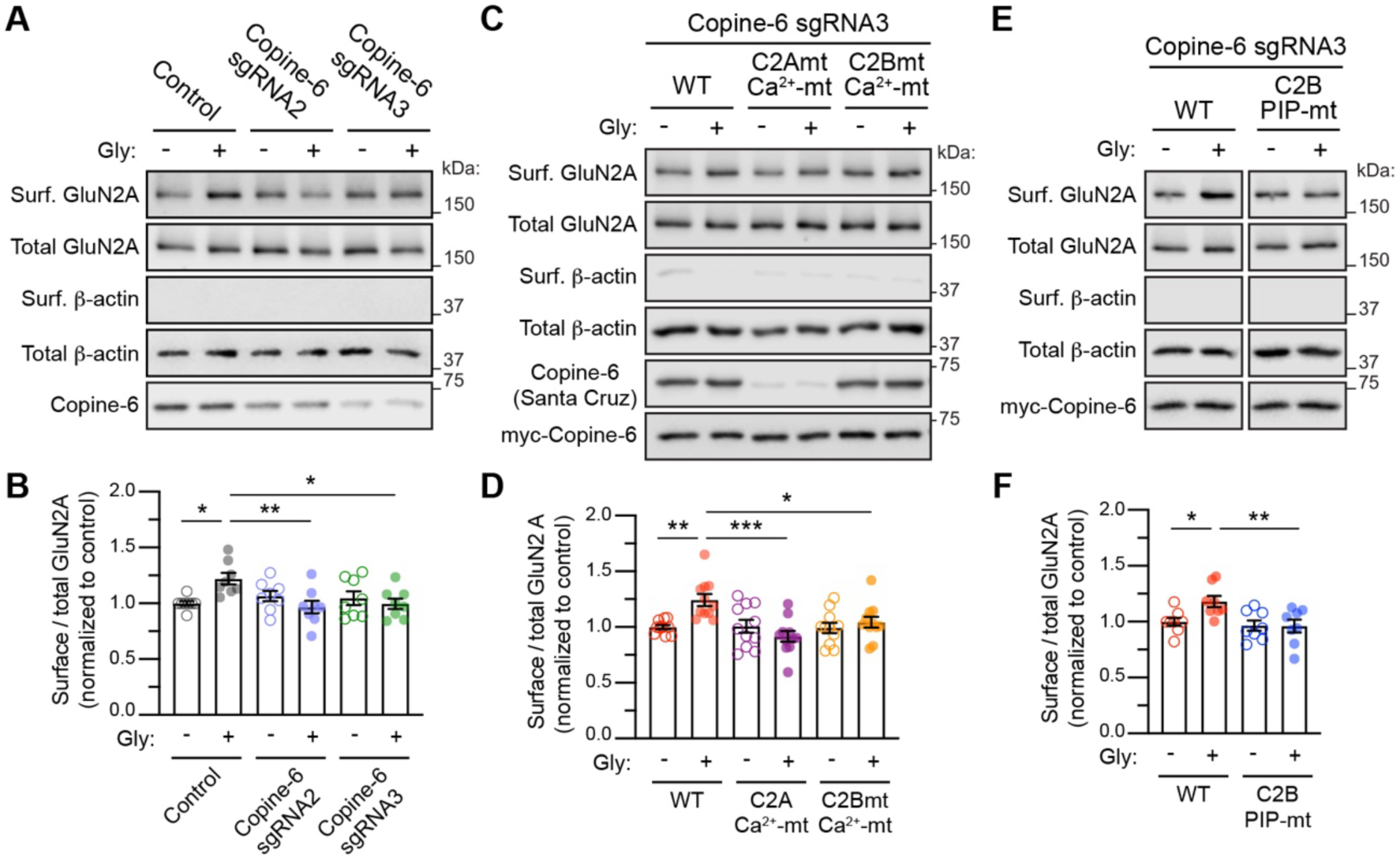
Copine-6 is required for activity-induced GluN2A-NMDAR exocytosis. (A) Primary cortical neurons were transduced with lentiviral particles expressing Cas9-GFP, either alone (Control) or with Copine-6 sgRNA2 or sgRNA3 at DIV 9. At DIV15, transduced neurons were stimulated with glycine for 5 min and subjected to a surface biotinylation assay. The relative amounts of surface and total proteins were assessed by western blotting using specific antibodies against GluN2A, Copine-6 and β-actin. (B) Quantification of the surface to total GluN2A ratio in control and Copine-6 knockdown neurons. Data were normalized to unstimulated Cas9-GFP control neurons (*n* = 9 cultures per group from 7 independent experiments). (C and D) Surface biotinylation assays and quantifications of surface to total GluN2A ratio in Copine-6 knockdown neurons (Cas9-GFP-Copine-6 sgRNA3) expressing sgRNA3-resistant myc-Copine-6, either wild-type (WT), D93N/E95A (C2A Ca^2+^-mt) or D229/231/237N (C2B Ca^2+^-mt) Ca^2+^-binding mutants. (E and F) Surface biotinylation assays and quantifications of surface to total GluN2A ratio in Copine-6 knockdown neurons (Cas9-GFP-Copine-6 sgRNA3) expressing sgRNA3-resistant myc-Copine-6, either wild-type (WT) or R192Q/K197A (C2B PIP-mt) phospholipid-binding mutants. All data are represented as mean ± SEM. *n* = 8-11 cultures per group from 6 independent experiments. \**P* < 0.05, \*\**P* < 0.01 and \*\*\**P* < 0.001 using one-way ANOVA with a Dunnett’s multiple comparison test.

**Figure 8.**
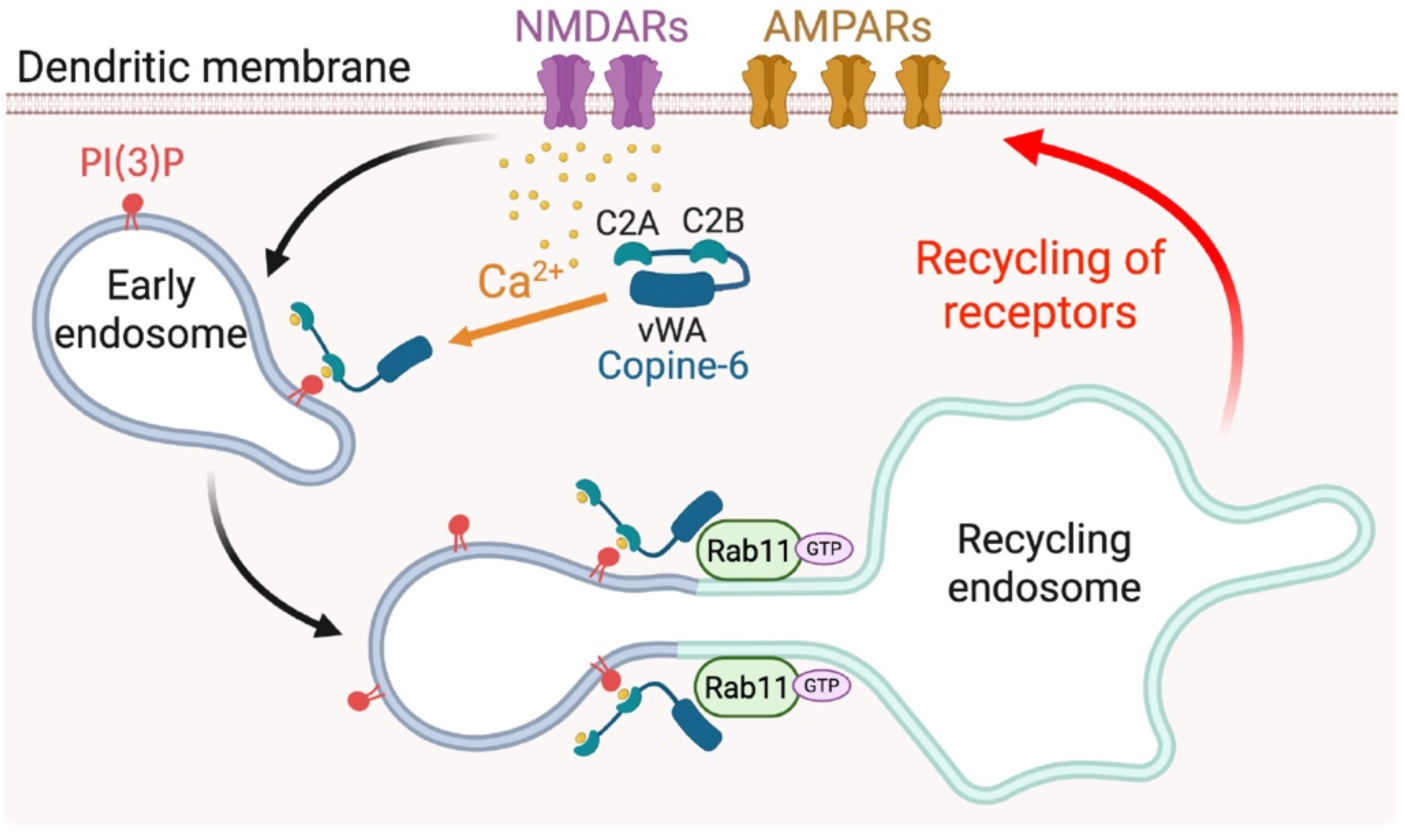
A proposed working model for the role of Copine-6 in regulating intracellular membrane dynamics during synaptic potentiation. During LTP, the activation of NMDAR leads to an influx of Ca^2+^ into the postsynaptic compartment and activates downstream signaling pathways that enhance the synthesis of PI(3)P in early endosomes and increase the level of active Rab11 at the recycling endosomes. The elevation of postsynaptic Ca^2+^ level is also sensed by Copine-6, which facilitates the accumulation of Copine-6 in the early endosomes through PI(3)P binding. Active Rab11 recruits Copine-6 through its vWA domain to the recycling endosomes, effectively coupling the early and recycling endosomes and facilitate the forward trafficking of cargo molecules, including AMPA and NMDA receptors towards the plasma membrane. Image was created with Biorender.com.

## Discussion

The maintenance of LTP relies on the continuous supply of AMPARs from the intracellular compartments to the neuronal plasma membrane. In this study, we identify the neuronal-specific Ca^2+^ and phospholipid-binding protein Copine-6 as a critical scaffolding molecule that mediates activity-dependent coupling of early and recycling endosomes in neurons (Fig. 8). Upon NMDAR activation, Copine-6 senses an increase in the level of Ca^2+^ concentration in the postsynaptic compartment and becomes accumulated in the early and recycling endosomes (33). Activity-dependent accumulation of Copine-6 into these intracellular endosomal compartments relies on two distinct mechanisms. Enhanced association of Copine-6 with the early endosomes relies on Ca^2+^-dependent binding to PI(3)P via the C2B domain. In contrast, the recruitment of Copine-6 to recycling endosomes is mediated by active Rab11 binding to the vWA domain. We propose that Copine-6 acts as a key molecule that senses and integrates various NMDAR-dependent signaling, including the rise in intracellular Ca^2+^ concentration, the increase in PI(3)P production, and elevated levels of Rab11-GTP to coordinately promote the forward trafficking of AMPARs to the plasma membrane during synaptic potentiation.

Copine-6 has been shown to interact with PS in a Ca^2+^-dependent manner, thought to mediate its targeting to the plasma membrane (33, 35–38). PS is enriched on the inner leaflet of the plasma membrane, but an increase in intracellular Ca^2+^ concentration also triggers the translocation of Copine-6 to endosomal compartments in heterologous cells and neurons (33, 38). One of the major findings that Copine-6 also interacts with PIPs upon Ca^2+^ binding provides a molecular basis that explains the activity-dependent translocation of Copine-6 to the early endosomes in neurons. MD simulations and structural alignment analyses reveal an analogous mechanism of phospholipid binding among Copine-6, PKCα, and Syt-1, whereby PS and PIPs occupy the Ca^2+^-binding loops and the cationic β-groove, respectively (39). Although both C2A and C2B domains of Copine-6 bind PS, only the C2B domain can interact with PIPs. The diversity of binding modes and phospholipid selectivity between the two Copine-6 C2 domains may further enhance Copine-6 membrane targeting while providing spatial specificity for association with multiple membrane compartments in cells. Ca^2+^-dependent engagement of PS together with PIPs may provide a coincidence-detection mechanism that targets Copine-6 to specific membrane microdomains during neuronal activity, thereby positioning its C-terminal vWA domain to interact with other proteins involved in synaptic plasticity.

Mutations of key positively charged residues (R192Q and K197A) in the C2B domain were initially designed to abolish Copine-6 binding to both PS and PIPs. The C2B PIP-mt robustly inhibits Ca^2+^-dependent binding to PI(3)P, PI(5)P, and to a lesser extent, PI(4)P; however, it does not affect Copine-6 binding to PS. Nevertheless, this mutant allows us to probe the functional significance of the Copine-6-PIP interaction in the activity-induced translocation of Copine-6 to early endosomes. Interestingly, the C2B PIP-mt does not affect the targeting of Copine-6 to these FYVE-mCherry-enriched compartments under basal conditions, suggesting that Copine-6 association with endosomes can be mediated by another mechanism that is independent of PIP binding. One potential candidate is the vWA domain, which was previously shown to mediate Copine-6 association with transferrin-positive vesicles in heterologous cells (38). We propose that the newly synthesized PI(3)P in early endosomes (27) actively recruits cytosolic, Ca^2+^-bound Copine-6 to promote the recycling of cargo molecules during synaptic potentiation.

An interesting feature of the Copine-6 C2B PIP-mt is its enhanced association with the Rab11-positive recycling endosomes. PI(4)P, which is enriched in the recycling endosomes, is unlikely to mediate this interaction due to its reduced interaction with the C2B PIP-mt. Instead, we found that Rab11 can interact with Copine-6, primarily through its vWA domain. The constitutively active Rab11 S20V binds Copine-6 with higher affinity, indicating that Copine-6 is a downstream effector of active, GTP-bound Rab11. Consistent with a previous study, the interactions of Copine-6 with the small GTPases, Rac1 and Rab11, are independent of its association with Ca^2+^ (35). Our data suggest that activity-induced translocation of Copine-6 to early and recycling endosomes occurs in parallel, via Ca^2+^-bound C2B-PI(3)P binding and GTP-bound Rab11-vWA interaction, respectively. The interaction between Copine-6 and small GTPases may represent a common molecular module that expands the functional repertoire of Copine-6 in regulating intracellular membrane dynamics and membrane trafficking.

Efficient fusion of recycling endosomes to the neuronal plasma membrane is essential for supplying surface AMPARs during LTP (11, 13–15, 29, 30, 43, 45). Prior to that, AMPARs must be sorted from early endosomes and trafficked to recycling endosomes. Our study identified Copine-6 as a molecular bridge that promotes the coupling of EEA1-positive early endosomes with Rab11-positive recycling endosomes, in a process that relies on Copine-6 binding to Ca^2+^ and PI(3)P. Molecular manipulation of Copine-6 protein expression, Ca^2+^, and PIP bindings effectively blocks glycine-induced insertion of AMPARs to the neuronal plasma membrane. The other known molecule that fulfills a similar role is the Rab4 effector, GRASP1, which is essential for AMPAR recycling during LTP (31, 32). Mechanistically, GRASP1 segregates Rab4 from Rab5-positive membranes by interacting with syntaxin-13, which is enriched in the Rab11-positive recycling endosomes (31). GRASP1 is also a GRIP1-interacting partner that directly binds to the C-terminal tail of the GluA2 subunit of AMPARs (52). It is worth noting that while Rab4 supplies membranes during structural plasticity, one study has shown that it is not required for AMPAR transport to the plasma membrane during LTP (30). Despite this, our findings suggest that during LTP, multiple mechanisms exist to coordinate the flow of intracellular AMPAR-containing vesicles/endosomes towards recycling compartments.

While the contribution of dendritic membrane trafficking to the maintenance of LTP is clear, it likely also delivers other important membrane proteins, such as TrkB (53) and GluN2A-NMDARs (46–51), in addition to AMPARs (45). Here, we found that Copine-6 is also responsible for activity-induced insertion of GluN2A-containing NMDARs into the plasma membrane during synaptic potentiation. The growing list of cargo molecules, including AMPARs (33), NMDARs (this study), TrkB (34), and TRPM3 (54), that interact with Copine-6 indicates its broader role as an essential Ca^2+^ sensor regulating activity-dependent membrane trafficking in neurons, thereby supporting both functional and structural plasticity (33–35). Future investigation aimed at identifying the composition of Copine-6-associated endosomes/vesicles is warranted. In conclusion, our findings offer a mechanistic framework for understanding how transient Ca^2+^ elevations in the postsynaptic compartments are integrated and translated into localized signaling events that coordinate endosomal membrane dynamics and forward trafficking of plasticity-related cargo molecules required to maintain LTP.

## Materials and Methods

### Animals

Adult female Sprague-Dawley rats and their embryos (male and female) at embryonic day 18 were used to isolate primary hippocampal and cortical neurons. All research procedures involving the use of animals were conducted in accordance with the Australian Code of Practice for the care and use of animals for scientific purposes and were approved by the University of Queensland Animal Ethics Committee (2021/AE511).

### Antibodies

The following antibodies were obtained commercially: rabbit anti-GluA1 C-terminal (Cat# ab1504, Millipore), mouse anti-GluA1 C-terminal (Cat# 182011, Synaptic Systems), mouse anti-β-actin (Cat# sc-47778, Santa Cruz Biotechnology), rabbit anti-GluN2A (Cat# 04-901, Millipore), rabbit anti-Copine-6 (Cat# 13782-1-AP, Proteintech), mouse anti-Copine-6 (Cat# sc-136357, Santa Cruz Biotechnology), mouse anti-EEA1 (Cat# 610456, BD Biosciences), chicken anti-GFP (Cat# GFP-1020, Aves Labs), mouse anti-GST (Cat# 66001-2-Ig, Proteintech), mouse anti-myc (Cat# sc-40, Santa Cruz Biotechnology) and mouse anti-Rab11 (Cat# 610656, BD Biosciences). Alexa-conjugated secondary antibodies were purchased from Thermo Scientific. HRP-conjugated sheep anti-mouse and donkey anti-rabbit secondary antibodies were obtained from GE Healthcare, whereas HRP-conjugated goat anti-chicken secondary antibodies were from Santa Cruz Biotechnology.

### DNA constructs

Various plasmids that encode myc-Copine-6, HA-Copine-6, Copine-6-GFP, GST-Copine-6, His-Copine-6, Copine-6 sgRNA (5’-GAGCCTCTCGAGTAGAGCTG-3’) and GST-GluN2A (residues 1213-1464) have been described previously (33, 46). Copine-6 C2B phospholipid-binding mutant (R192Q/K197A) was generated by an overlapping PCR protocol with the following primers: 5’-TGG CAG ACT GAG GTG GTG GCA AAC AAT TTG AAC CCC AGC TGG-3’ (sense) and 5’-GTT TGC CAC CAC CTC AGT CTG CCA GAC CAG TTG GTC ACT CTG-3’ (anti-sense). pCAG-SEP-GluA1 and pEGFP-Rab11a were gifts from Prof. Richard Huganir (Johns Hopkins University) and Prof. Jenny Stow (University of Queensland), respectively.

### Primary neuronal cultures

Primary rat hippocampal and cortical neurons were prepared from Sprague-Dawley rat embryos at embryonic day 18 as previously described (19, 33). In brief, cortical and hippocampal tissues were digested in 30 U papain (Worthington) solution for 20 min at 37°C, then triturated with fire-polished Pasteur pipettes to obtain a single-cell suspension. Neurons were plated on poly-L-lysine-coated 12-well plates at densities of 80,000 (hippocampal) and 200,000 (cortical) cells per well. Cultured neurons were maintained in Neurobasal medium supplemented with 2% B-27 (Invitrogen), 2 mM GlutaMAX (Invitrogen) and 1% penicillin/streptomycin (Invitrogen) in a 37°C incubator with 5% CO_2_. Neurons were fed twice weekly with Neurobasal medium containing 0% (hippocampal) or 1% FBS (cortical). To inhibit glial proliferation in cortical cultures, 5 μM uridine and 5 μM 5’-fluoro-2’-deoxyuridine (Sigma) were added to cortical cultures at days *in vitro* (DIV) 5. Hippocampal neurons were transfected at DIV 12-13 using Lipofectamine 2000 (Invitrogen) per the manufacturer’s instructions and processed at DIV 15-16.

### Glycine-induced chemical LTP (cLTP)

Mature primary neurons at DIV 15-16 were washed and incubated in pre-warmed artificial cerebrospinal fluid with low Mg^2+^ (ACSF; containing 0.4 mM MgCl_2_, 2 mM CaCl_2_, 120 mM NaCl, 5 mM KCl, 30 mM glucose, 25mM HEPES, pH 7.4) supplemented with 20 µM bicuculline (Abcam), 5 µM strychnine (Sigma) and 0.5 µM tetrodotoxin (Abcam) for 1 h at 37°C. cLTP was induced by incubating neurons in pre-warmed low Mg^2+^ ACSF containing 200 µM glycine, 20 µM bicuculline and 5 µM strychnine for 5 min at room temperature.

### Lentivirus packaging and transduction

Lentiviral particles were produced in HEK293T cells by transfecting them with 7 μg of the target plasmid, 3 μg each of pMD2.G (Addgene #12259), pRSV-Rev (Addgene #12253) and pMDLg/pRRE (Addgene #12251) plasmids using calcium-phosphate precipitation. The supernatant containing lentivirus was collected 48 h post-transfection, filtered through a 0.45 μm cellulose acetate membrane with low protein binding, and concentrated either by ultracentrifugation at 106,559 *g* for 2 h at 4°C in a Beckman SW 32 Ti rotor or using PEG-it Virus Precipitation Solution (System Biosciences) per the manufacturer’s instructions. The viral pellet was resuspended in Neurobasal medium, snap-frozen in liquid nitrogen, and stored at -80°C. Cortical neurons were transduced with lentiviral particles at DIV 8-9 (overnight) for Copine-6 sgRNA-knockdown and again at DIV 12 (6 h) for Copine-6 overexpression (rescue experiments), followed by incubation for an additional 3 days before analysis.

### Immunocytochemistry

Primary hippocampal neurons were washed and fixed with Parafix solution (4% PFA and 4% sucrose in PBS) for 15 min, permeabilized (0.25% Triton-X100) for 10 min and blocked (10% normal goat serum in PBS) for 1 h. Neurons were incubated with primary antibodies diluted in Can Get Signal® Immunoreaction Enhancer Solution (Cat# NKB-601, Toyobo Life Science) overnight at room temperature. Protein localization was visualized by staining with Alexa-conjugated secondary antibodies for 1 h. Images of neurons were collected using a Zeiss Plan Apochromat 63x/1.4 NA oil-immersion objective on an LSM710 confocal laser-scanning microscope. To quantify the colocalization of Copine-6 and intracellular endosomal markers, a region of interest was drawn on either the soma or the secondary dendrite to obtain the Manders’ coefficient using Just Another Colocalization Plugin (JACoP) in ImageJ software. The coefficient of variation (CV) for GFP-Rab11a in the dendrites was calculated by dividing the standard deviation by the mean GFP-Rab11a intensity along a 150-pixel distance secondary dendrite. The average CV for 4-5 dendrites per neuron was obtained.

### Proximity ligation assay (PLA)

Following cLTP, the association between endogenous GluA1 with exogenously expressed sgRNA-resistant HA-Copine-6 (wild-type or C2B PIP-mt) in Copine-6-depleted neurons was detected using the Duolink in Situ PLA kit (Sigma) according to the manufacturer’s protocol. Briefly, neurons were fixed, permeabilized, blocked and incubated with mouse anti-GluA1 (1:200) and rabbit anti-Copine-6 (1:150) antibodies overnight at room temperature. Neurons were incubated with mouse PLUS and rabbit MINUS Duolink PLA probes for 1 h at 37°C. The ligation reaction was then carried out for 30 min at 37°C. The rolling-circle amplification reaction for fluorophore labeling was performed at 37°C for 100 min. The number of PLA puncta per soma and 50 μm segment of a primary dendrite was counted and normalized to control cells.

### SEP-GluA1 insertion assay

Plasma membrane insertion of SEP-GluA1 in living hippocampal neurons was visualized on a Zeiss ELYRA microscope under the TIRF mode with a 100X oil-immersion objective. To visualize newly inserted AMPARs on the dendritic membranes, pre-existing surface SEP-GluA1 was photobleached with 100% laser power for 10 s before data acquisition. Images were acquired at 0.5 Hz for 5 min. Signals lasting at least 4 frames (8 s) were manually scored as insertion events from the *y*-t rendered images. The *y*-t rendering was performed in ImageJ software as described previously (33). Data were expressed as the total number of SEP-GluA1 insertions per μm of dendrite over 5 min. The typical length of secondary dendritic segments analyzed was 35 to 50 μm.

### Surface biotinylation assay

The surface biotinylation assay was performed to measure the levels of endogenous GluA1 and GluN2A at the plasma membrane. Live neurons were rinsed with cold ACSF (pH 8.2) and incubated with ACSF (pH 8.2) containing 0.5 mg/ml sulfo-NHS-SS-Biotin (CovaChem) for 30 min at 4°C. Free biotin was quenched by washing cells three times with ice-cold Tris-buffered saline (TBS). Neurons were then lysed in RIPA buffer (1% Triton X-100, 0.5% Na-deoxycholate, 0.1% SDS, 2 mM EDTA, 2 mM EGTA, 50 mM NaF, 10 mM Na-pyrophosphate, 150 mM NaCl, 50 mM Tris pH 7.4) supplemented with EDTA-free protease inhibitor cocktail (Roche) for 30 min at 4°C. Lysates were cleared by centrifugation at 20,627 *g* at 4°C for 20 min. A small fraction of cleared lysates was collected as input to determine the total GluA1 or GluN2A levels, and the remaining lysates were incubated with Neutravidin beads (Thermo Fischer Scientific) overnight at 4°C to isolate biotinylated surface proteins. The beads were washed three times with ice-cold RIPA buffer, and bound proteins were eluted with 2X SDS sample buffer and heated at 50°C for 30 min. Samples were analyzed by western blotting. The ratio of surface/total GluA1 and surface/total GluN2A of each sample was normalized to the non-stimulated control groups.

### GST pull-down assay

HEK293T cells were cultured in DMEM containing 4.5 g/L glucose, 10% FBS, 50 U/mL penicillin, and 50 mg/mL streptomycin in a humidified 5% CO_2_ incubator at 37°C. Cells were transfected using the calcium precipitation method and lysed in ice-cold RIPA buffer containing EDTA-free protease inhibitor cocktail. For the myc-Copine-6 and GST-GluN2A-C-tail pull-down, cells were lysed with ice-cold cell lysis buffer (1% Triton X-100, 1 mM EDTA, 1 mM EGTA, 50 mM NaF, 5 mM Na-pyrophosphate in PBS) plus protease inhibitors. Lysates were cleared by centrifugation at 20,627 *g*, 4°C for 20 min. A small amount of cleared lysate was collected for input, and the remaining lysate was incubated with glutathione agarose beads (Thermo Scientific) overnight at 4°C. Beads were washed four times with ice-cold RIPA or cell lysis buffer, and bound proteins were eluted with 2X SDS sample buffer at 100°C for 10 min. Samples were analyzed by western blotting.

### Purification of recombinant Copine-6 protein

Plasmids encoding His-Copine-6 (either full-length or individual C2 domains) were transformed into *E. coli* BL21(DE3) cells and cultured in LB broth at 37°C until the density reached an OD of 0.8. Recombinant protein expression was induced by adding 0.5 mM isopropyl 1-thio-β-D-galactopyranoside (IPTG) to the culture medium, and the cultures were incubated at 20°C overnight. Cells were harvested by centrifugation at 6,000 *g* for 10 min at 4°C. Cell pellets were resuspended in ice-cold buffer (50 mM Tris, 300 mM NaCl, 5% glycerol, 10 mg/mL DNase I, 100 μM PMSF, 1 mM β-mercaptoethanol, pH 8.0) and lysed by mechanical disruption at 30 kpsi using a Constant Systems cell disrupter. The lysate was cleared by centrifugation at 50,000 *g* for 30 min at 4°C. Proteins were purified by affinity chromatography using Ni^2+^ Sepharose protein purification resin (Cytiva) on a gravity column. Proteins were eluted in 50 mM Tris, 300 mM NaCl, 5% glycerol, 250 mM imidazole, 1 mM β-mercaptoethanol, pH 8.0. The affinity-purified proteins were then subjected to size-exclusion chromatography (Superdex-200 16/600 HiLoad column) on an AKTA pure (GE Healthcare) with a buffer containing 50 mM HEPES, 200 mM NaCl, 0.5 mM TCEP, pH 7.5. The eluted protein was concentrated using a Centricon Ultra-10 kDa centrifugal filter (Millipore) and analyzed by the Bradford assay (BioRad) to determine protein concentration.

### Liposome pelleting assay

The control liposome (PC/PE) was prepared with a 90:10 molar ratio of POPC (1-palmitoyl-2-oleoyl-sn-glycero-3-phosphocholine): POPE (1-palmitoyl-2-oleoyl-sn-glycero-3-phosphoethanolamine). Liposome containing PS was prepared in a molar ratio of 60% POPC, 10% POPE and 30% POPS (1-hexadecanoyl-2-(9Z-octadecenoyl)-sn-glycero-3-phospho-L-serine). Liposome containing PIPs was prepared in a molar ratio of 80% POPC, 10% POPE and 10% of either PI(3)P [1,2-dioctanoyl-sn-glycero-3-(phosphoinositol-3-phosphate)], PI(4)P [1,2-dioleoyl-sn-glycero-3-phospho-(1′-myo-inositol-4′-phosphate)] or PI(5)P [1,2-dioleoyl-sn-glycero-3-phospho-(1′-myo-inositol-5′-phosphate)]. All liposomes were purchased from Avanti Polar Lipids and freshly prepared in a total volume of 500 μL of chloroform. Lipid films were generated by evaporating the solvent using a round-bottom glass flask under a nitrogen gas stream, followed by overnight incubation in a vacuum desiccator. Multilamellar vesicles (MLVs) were generated by hydrating the lipid film with 500 μL sucrose solution (25 mM Tris, 220 mM sucrose, pH 7.4), followed by agitation for 1 h and 10 rapid freeze-thaw cycles. MLV suspensions were pelleted at 180, 000 *g* for 30 min at 4°C by ultracentrifugation (TLA-55 rotor, Optima TL Ultracentrifuge, Beckman Coulter). The MLV pellet was resuspended in a buffer containing 50 mM HEPES, 200 mM NaCl, 0.5 mM TCEP, pH 7.5. The liposome binding assay was performed in a 50 μL reaction comprising 10 μM of recombinant Copine-6 protein, 25 μL of liposomes, with or without 2 mM CaCl_2_. The reaction mixture was incubated for 30 min at room temperature, followed by ultracentrifugation at 400,000 *g* (TLA-100 rotor, Optima TL Ultracentrifuge, Beckman Coulter) for 30 min at 4°C. The supernatant and liposome pellet fractions were analyzed by SDS-PAGE and Coomassie Blue staining.

### Lipid-strip binding assay

A PIP Strips membrane (Invitrogen) was blocked in 3% bovine serum albumin (BSA, fatty acid free, Sigma) in TBS (containing 0.1% Tween-20, TBS-T) for 1 h at room temperature. Purified recombinant GST-Copine-6 (residues 1-305, 0.5 μg/ml) protein was then added and incubated for another hour. After washing five times with TBS-T solution, bound Copine-6 was detected by western blotting using anti-GST antibodies.

### Western blotting

Samples were loaded in 7.5%, 10% or 12% SDS-PAGE gels and separated at 110 V for 1–2 h. Proteins were then transferred to a PVDF membrane at 100 V for 2 h. Membranes were blocked in 5% skim milk TBS-T for 1 h and washed in TBS-T three times at 5 min intervals before overnight incubation with primary antibodies at 4°C. Membranes were washed five times in 1% milk/TBS-T and incubated with HRP-conjugated secondary antibodies (GE Healthcare, 1:10,000) for 1 h at room temperature. They were washed extensively and developed using the enhanced chemiluminescent method (PerkinElmer). Images were acquired with a LiCOR imaging system and quantified with ImageStudio software.

### Molecular dynamics (MD) simulation

The AlphaFold-predicted structure of the C2B domain of Copine-6 (Uniprot ID: D4ACG7) was used to investigate its dynamics in the presence of PS, PI(3)P, and Ca²⁺ ions. The model (AF-D4ACG7-F1-model_v4.pdb) was of high confidence over the simulated domain (median per-residue pLDDT 94.0; 95% of residues above 70 and 64% above 90), including all lipid- and Ca²⁺-coordinating residues (pLDDT 87.5–97.2 for D167, D173, Y179, R192, K197, Y228, D229, Y230, D231 and D237). The only residues below 70 were the four at the N-terminal truncation boundary (S148-N151) and three in the G183-Q188 loop. The initial locations of five Ca²⁺ ions were derived from the rat Syt-1 C2B domain structures [PDB IDs: 1UOV and 2YOA] (55, 56).

To simplify the simulations, only the headgroups of PS and PI(3)P (referred to as hPS and hPI3P, respectively) were modeled, while their fatty acid tails were represented by methyl groups. The topologies of hPS and hPI3P were generated using the R.E.D. Server (57) with the use of the quantum mechanics program Gaussian16_C.01 (58) using a two-stage RESP fit to electrostatic potentials computed at the HF/6-31G* level and converted to GROMACS format with AmberTools 2023 (59) and ACPYPE (60). The net charges of hPS and hPI3P are -1 and -7, respectively. The positions of hPS and hPI3P were based on their locations in the rat Syt1 C2B–PS [PDB ID: 2YOA] (56) and rat PKCα C2–PI(4,5)P₂ [PDB ID: 3GPE] (40) complexes, respectively.

All simulations were performed using AMBER99SB (61) and the SPC/E water model (62) on GROMACS version 2022.3 (63). Hydrogens were added with gmx pdb2gmx, which assigns standard protonation states at neutral pH. The system was solvated in a periodic cubic box with a 1 nm buffer between the complex and the box walls. Cl⁻ ions were added to neutralize the net charge of the system only, and no additional NaCl was included. Bond lengths were constrained using the LINCS algorithm (64), and dynamics were integrated with a 2 fs timestep using the leap-frog algorithm (65) with a Verlet cutoff (66) scheme and neighbor lists updated every 10 steps. Long-range electrostatics were treated using the particle-mesh Ewald method (67). Temperature was maintained at 300 K using the V-rescale thermostat with a coupling time of 0.1 ps (68), and pressure was controlled at 1 atm using the Parrinello–Rahman barostat with a pressure relaxation time of 2 ps (69). The cutoff distances for van der Waals and short-range electrostatic interactions were set to 1.0 nm.

Energy minimization was first performed using 50,000 steps of the steepest descent algorithm. The system was then equilibrated for 200 ps at constant volume, followed by 200 ps at a constant pressure of 1 atm. Harmonic positional restraints with a force constant of 1000 kJ mol⁻¹ nm⁻² were applied to the heavy atoms throughout equilibration. Production MD simulations were subsequently carried out for 100 ns at 300 K in the NPT ensemble in triplicate, with no restraints on the protein atoms. Initial velocities for each replicate were generated independently from a Maxwell– Boltzmann distribution at 300 K using a randomly selected seed (gen_vel = yes, gen_seed = -1), so the three replicates are statistically independent; all replicates started from the same initial complex geometry, and their divergence therefore reflects thermal sampling rather than differences in the starting pose. Trajectory visualization was performed using VMD (70). Distance analyses were conducted with in-house Python scripts.

### Statistical analysis

The sample size (*n*) reported in figure legends represents individual neurons or wells generated from at least three independent experiments, unless otherwise stated. Statistical analysis was performed in GraphPad Prism 9.0 using one-way analysis of variance (ANOVA) with Tukey’s or Dunnett’s post-hoc multiple comparisons tests. For comparison between two groups, a two-tailed unpaired t-test was employed. All data are reported as mean ± standard error of the mean (SEM).

## Acknowledgments

We thank R. Amor and the Queensland Brain Institute (QBI) microscopy team for support with microscopy. This work was supported by Australian National Health and Medical Research Council (NHMRC) Project Grant GNT1138452 (to V.A. and B.M.C.), Australian Research Council (ARC) Discovery Project Grant DP220101645 (to V.A.), Clem Jones Centre for Ageing Dementia Research Flagship Project Grant (to V.A. and J.W.), Donald & Joan Wilson Foundation Project Grant (to J.W.), and a generous donation from Mrs Kay Bryan OAM (to V.A.). V.A. holds an ARC Future Fellowship (FT220100485). B.M.C. and D.B.A. hold NHMRC Investigator Grants. S.E.J., A.B.B., L.Z., and G.G. were supported by University of Queensland Research Training Scholarships. Imaging was performed at the QBI Advanced Microscopy Facility, supported by the Australian Government through ARC LIEF grant LE130100078.

## Author Contributions

Conceptualization: J.Z.A.T., B.M.C. and V.A. Methodology: J.Z.A.T, A.B.B., M.C., T.B.N., K.C., S.W., D.B.A., B.M.C. and V.A. Investigation: J.Z.A.T, A.B.B., M.C., T.B.N., S.E.J., G.G., L.Z., K.C. and J.W. Visualization: J.Z.A.T., T.B.N., L.Z., and V.A. Funding acquisition: J.W., B.M.C. and V.A. Project administration: V.A. Supervision: D.B.A., J.W., B.M.C. and V.A. Writing – original draft: J.Z.A.T. and V.A. All authors reviewed and edited the manuscript.

## Supplementary Figures and Videos

**Supplementary Figure 1.**
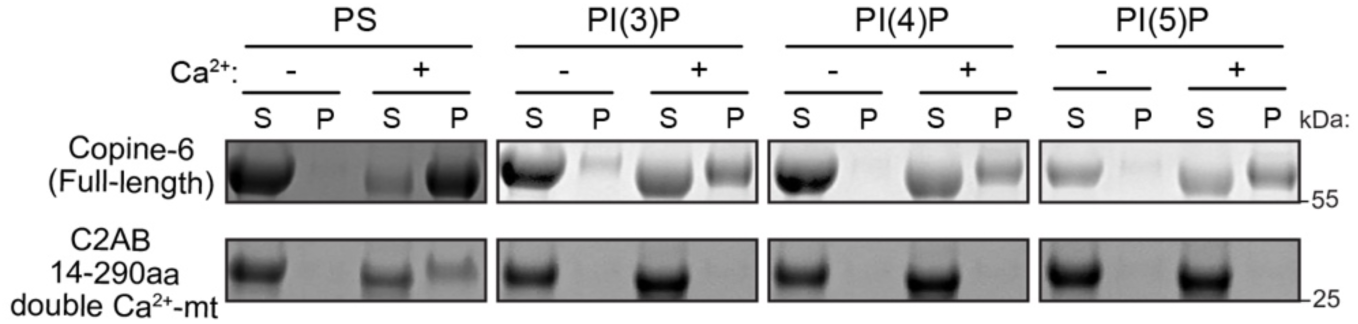
Copine-6 Ca^2+^-binding deficient mutant reduces phospholipid binding. Purified recombinant His-Copine-6 full-length proteins (1-557 amino acids) or His-Copine-6 (14-290 amino acids) Ca^2+^-binding mutant (D93N/E95A/D229N/D231N/D237N) were incubated with liposomes in the presence or absence of 2 mM Ca^2+^, followed by ultracentrifugation. The liposomes contain 0% (PC/PE), 30% PS or 10% PIPs. The unbound supernatant (S) and liposome-bound pellet (P) fractions were subjected to SDS-PAGE and Coomassie staining.

**Supplementary Figure 2.**
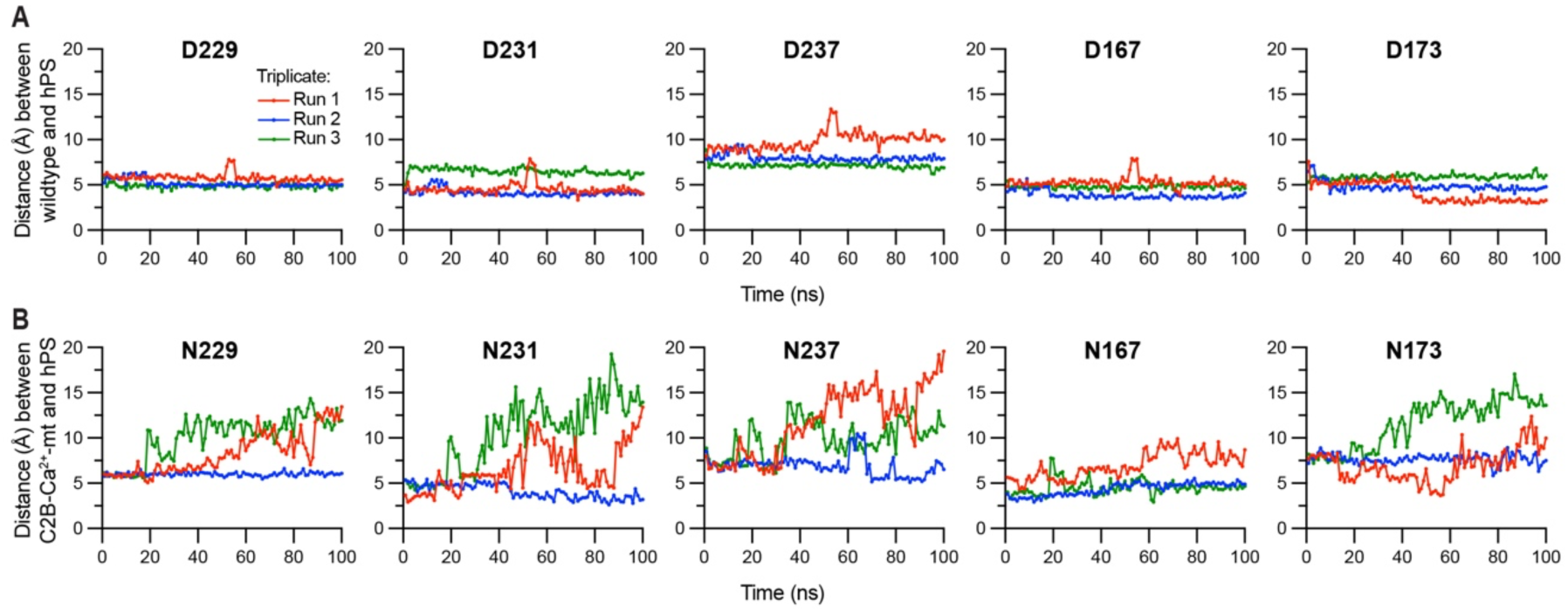
Stability of hPS interaction with Copine-6 Ca²⁺-binding residues. (A) The stability of hPS in the C2B domain of wild-type Copine-6 protein was evaluated based on the distances between hPS and the Asp residues (D167, D173, D229, D231, and D237) during simulation time. (B) The same analysis was also performed with Copine-6 C2B Ca^2+^-mt, in which the corresponding Asp residues were mutated to Asn. Each trace corresponds to data from an independent molecular dynamics (MD) simulation.

**Supplementary Figure 3.**
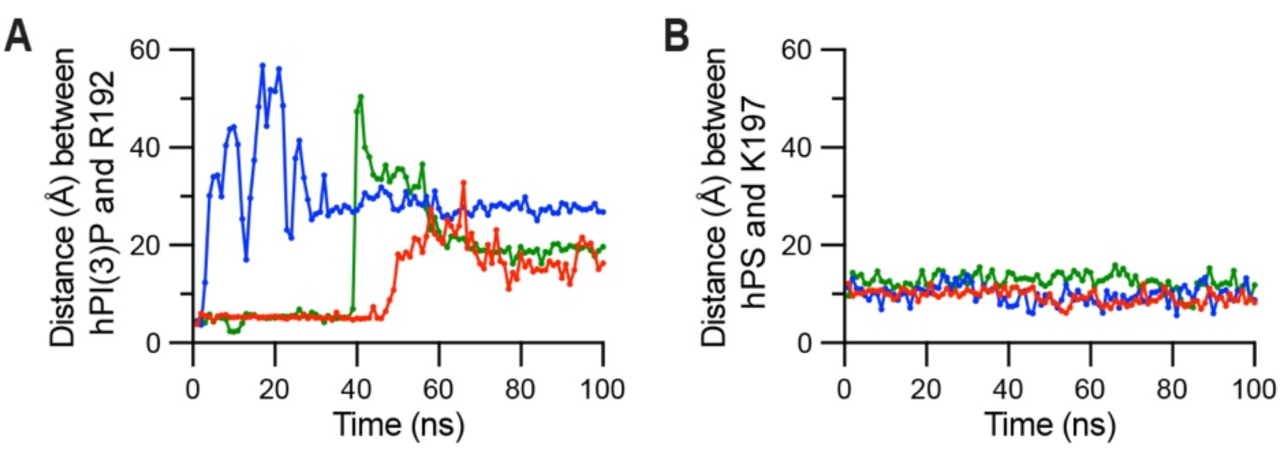
Stability of hPS and hPI(3)P with selected amino acids in the Copine-6 C2B domain. The stability of hPI(3)P and hPS in Copine-6 C2B domain was evaluated based on the distances between these phospholipid headgroups with Arg-192 (A) and Lys-197 (B), respectively, during simulation time. Each trace corresponds to data from an independent molecular dynamics (MD) simulation.

**Supplementary Figure 4.**
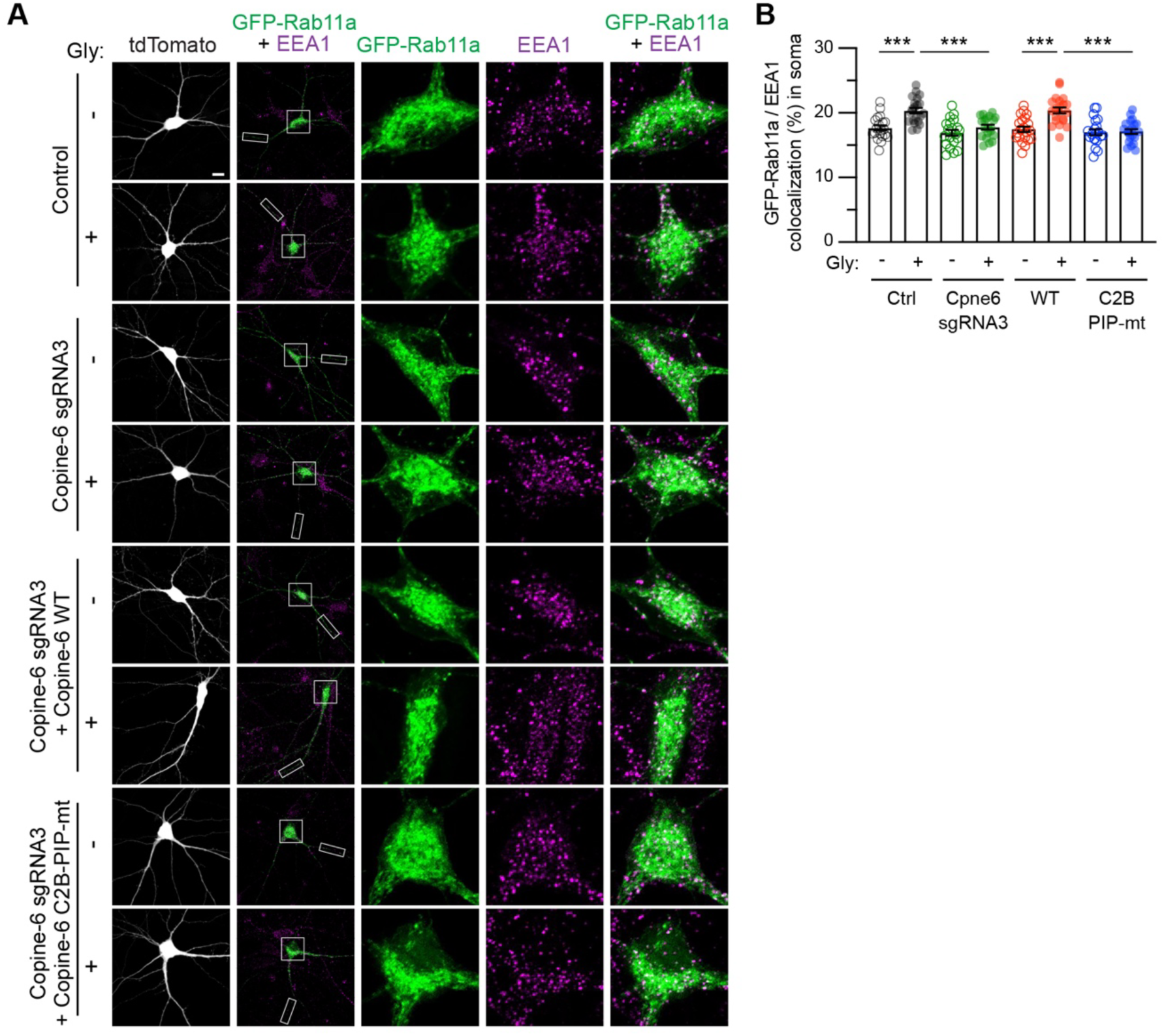
Copine-6 regulates the coupling between EEA1- and Rab11a-positive endosomes upon synaptic potentiation. (A) The plasmid encoding GFP-Rab11a (green) was transfected into primary hippocampal neurons expressing Cas9-T2A-tdTomato only (control), Cas9-T2A-tdTomato-Copine-6 sgRNA3 (Cpne6 knockdown), Cas9-T2A-tdTomato-Copine-6 sgRNA3 plus sgRNA3-resistant Copine-6 wild-type (Rescue WT) or Cas9-T2A-tdTomato-Copine-6 sgRNA3 plus sgRNA3-resistant Copine-6 R192Q/K197A (Rescue C2B PIP-mt) at DIV 12. Neurons were stimulated with glycine for 5 min at DIV15, fixed and immunostained for endogenous EEA1 (magenta). Representative images of transfected neurons and their soma (inset) are shown. Scale bars, 10 μm. (B) Quantification of GFP-Rab11a in EEA1-positive compartments in the soma using Mander’s coefficient (n = 20-21 neurons from 3 independent experiments). All data are represented as mean ± SEM. *** P < 0.001 using one-way ANOVA with Tukey’s multiple comparison test.

**Supplementary Figure 5.**
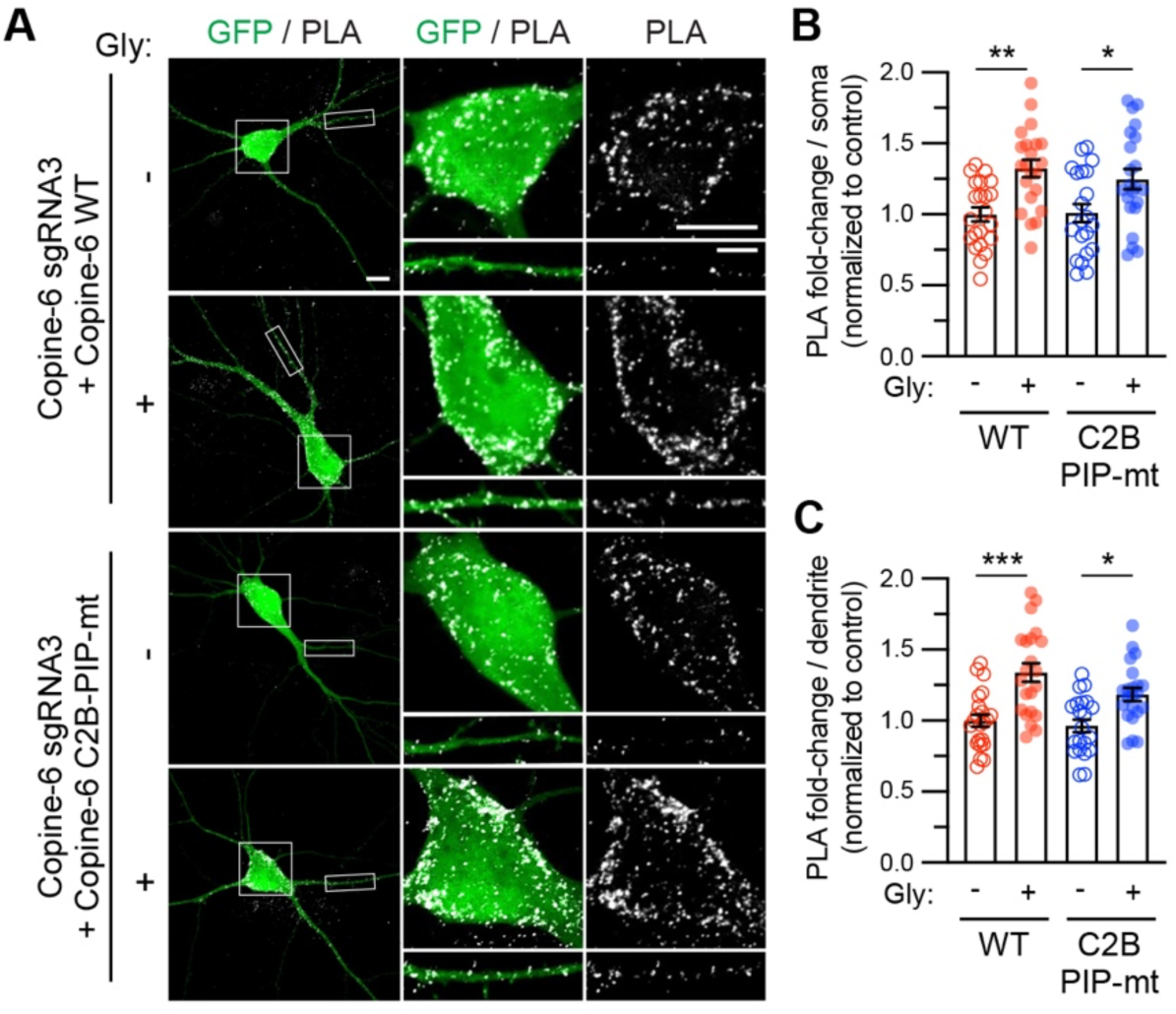
Copine-6 and GluA1 proximal interaction does not depend on Copine-6 binding to phospholipids. (A) Primary hippocampal neurons were co-transfected with plasmids encoding Cas9-GFP and Copine-6 sgRNA3, together with myc-Copine-6 sgRNA3-resistant constructs, either wild-type or C2B PIP-mt at DIV 12. At DIV 15, neurons were stimulated with glycine for 5 min and subjected to a proximity ligation assay (PLA) with GluA1 and Copine-6 antibodies. Representative images showing the PLA signals (grey) in the soma and dendrites. Scale bar, 10 μm. (B and C) Quantification of the number of PLA puncta in the (B) soma and (C) dendrites. Data are normalized to the unstimulated myc-Copine-6 WT group (n = 21-22 neurons per group from 4 independent experiments). All data are represented as mean ± SEM. *P < 0.05; **P < 0.01; ***P < 0.001 using one-way ANOVA with a Tukey’s multiple comparison test.

**Supplementary Figure 6.**
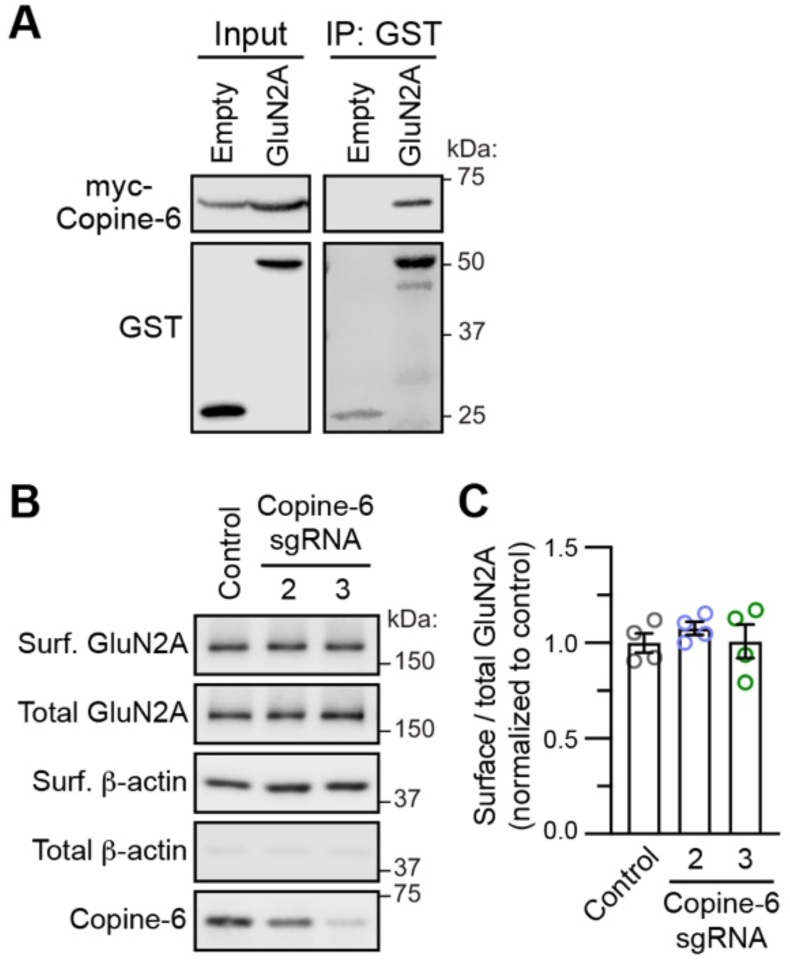
Copine-6 is not required for the steady-state expression of GluN2A-NMDARs under basal conditions. (A) GST pull-down experiments from HEK293T cells co-expressing Myc-Copine-6 with GST alone or GST-GluN2A carboxyl-tail (1213-1464 amino acids). Total lysates and bound proteins were immunoblotted with anti-Myc and anti-GST antibodies. (B) Primary cortical neurons were transduced with lentiviral particles expressing Cas9-GFP, either alone or with Copine-6 sgRNA2 or sgRNA3 at DIV 9. Transduced neurons were subjected to a surface biotinylation assay at DIV 15. The relative amounts of surface and total proteins were assessed by western blotting using specific antibodies against GluN2A, Copine-6 and β-actin. (C) Quantification of the surface/total GluN2A ratios in Copine-6-depleted neurons, normalized to control neurons (*n* = 4 cultures from 2 independent experiments). All data represent mean ± SEM.

**Supplementary Video 1. Molecular dynamics simulation of the wild-type Copine-6 C2B domain interacting with PS and PI(3)P headgroups.** Representative trajectory from a 100 ns molecular dynamics simulation of the AlphaFold-predicted Copine-6 C2B domain in the presence of dicaproyl phosphatidylserine (DCPS) and PI(3)P headgroups and Ca²⁺ ions. The Copine-6 C2B domain is shown as a cartoon, PS and PI(3)P headgroups as stick representations, and Ca²⁺ ions as blue spheres.

**Supplementary Video 2. Molecular dynamics simulation of the Ca2+-binding deficient Copine-6 C2B mutant interacting with PS and PI(3)P headgroups.** Representative trajectory from a 100 ns molecular dynamics simulation of the AlphaFold-predicted Copine-6 C2B Ca^2+^-mt in the presence of dicaproyl phosphatidylserine (DCPS) and PI(3)P headgroups and Ca²⁺ ions. The Copine-6 C2B Ca^2+^-mt is shown as a cartoon, PS and PI(3)P headgroups as stick representations, and Ca²⁺ ions as blue spheres.

